# Bayesian inference of admixture graphs on Native American and Arctic populations

**DOI:** 10.1101/2022.09.06.506725

**Authors:** Svend V Nielsen, Andrew H. Vaughn, Kalle Leppälä, Michael J. Landis, Thomas Mailund, Rasmus Nielsen

## Abstract

Admixture graphs are mathematical structures that describe the ancestry of populations in terms of divergence and merging (admixing) of ancestral populations as a graph. An admixture graph consists of a graph topology, branch lengths, and admixture proportions. The branch lengths and admixture proportions can be estimated using numerous numerical optimization methods, but inferring the topology involves a combinatorial search for which no polynomial algorithm is known. In this paper, we present a reversible jump MCMC algorithm for sampling high-probability admixture graphs and show that this approach works well both as a heuristic search for a single best-fitting graph and for summarizing shared features extracted from posterior samples of graphs. We apply the method to 11 Native American and Siberian populations and exploit the shared structure of high-probability graphs to address the relationship between Saqqaq, Inuit, Koryaks, and Athabascans. Our analyses show that the Saqqaq is not a good proxy for the previously identified gene flow from Arctic people into the Na-Dene speaking Athabascans.

**Author Summary:** One way of summarizing historical relationships between genetic samples is by constructing an admixture graph. An admixture graph describes the demographic history of a set of populations as a directed acyclic graph representing population splits and mergers. The inference of admixture graphs is currently done via greedy search algorithms that may fail to find the global optimum. We here improve on these approaches by developing a novel MCMC sampling method, *AdmixtureBayes*, that can sample from the posterior distribution of admixture graphs. This enables an efficient search of the entire state space as well as the ability to report a level of confidence in the sampled graphs. We apply AdmixtureBayes to a set of Native American and Arctic genomes to reconstruct the demographic history of these populations and report posterior probabilities of specific admixture events. While some previous studies have identified the ancient Saqqaq culture as a source of introgression into Athabascans, we instead find that it is the Siberian Koryak population, not the Saqqaq, that serves as the best proxy for gene flow into Athabascans.

## Introduction

Admixture graphs^1^ provide a concise description of the historical demographic relationships between genetic samples of populations, assuming their relationships are the product of discrete, instantaneous splits and admixture events. The assumption of discrete, instantaneous events is clearly an oversimplification for most real data, but it facilitates interpretation and makes admixture graphs a popular first step in analyses. Each graph topology is associated with parameters capturing population divergence and admixture proportions, and once these are fitted to genetic data, the goodness of fit can be measured to determine how accurately the graph captures the historical relationship between samples. Inferring graph topologies, however, involves a combinatorial search, and since the space of graphs grows super-exponentially in the number of populations and the number of admixture events, an exhaustive search is typically not possible. Instead, the search for well-fitting topologies is often done manually or through greedy algorithms.

The most popular methods for estimating admixture graphs are *TreeMix* by Pickrell and Pritchard^2^ and *qpGraph* by Patterson et al.^1^, both of which take a greedy approach to searching the state space of graph topologies. qpGraph allows users to sequentially identify the best phylogenetic position of a possibly admixed population in a previously established admixture graph and evaluate the improved fit in terms of simple allele-sharing statistics. The program *MixMapper* by Lipson et al.^3^ employs a similar strategy and has options for fitting up to two admixture events simultanously. TreeMix estimates an admixture graph *de novo* by automatically estimating the best tree without admixture events followed by automatic, sequential insertion of the admixture branches.

In contrast to MixMapper and qpGraph, TreeMix searches through many potential admixture graphs without user input. To penalize deviations from the expected and observed allele sharing statistics, all three methods use a Gaussian model for the distribution of allele frequencies among populations. The implicit assumption in the Gaussian model is that changes in allele frequency due to genetic drift can be approximated as a Brownian motion process. This assumption dates back to the early work by Edwards and Cavalli-Sforza ^4^ and has recently re-emerged as a computationally attractive alternative to the full Wright-Fisher process. It has previously been used in several other methods aimed at modeling the joint distribution of allele frequencies among populations^5^.

There are also phylogenetic network methods that infer admixture graphs using sets of locus-specific gene trees as nuisance parameters which are either pre-estimated^6^ or integrated out using MCMC^7 8^. These approaches must evaluate the likelihood of each gene tree separately, making them more computationally expensive and therefore limited to fewer populations than the Gaussian drift models. To handle larger datasets, some methods summarize all the gene trees into a few statistics that are evaluated with a pseudo-likelihood^9 10^ for a small reduction in accuracy^10^. In terms of speed, these pseudo-likelihood methods are similar to the Gaussian drift methods. However, the Gaussian approach offers a way to compute a true likelihood, which we use in this paper.

The greedy search algorithms used by current methods do not guarantee that the inferred graph is optimal. In practice, the optimal graph found by a greedy search can potentially be very different from better-fitting, but never-discovered, graphs^11 12^. Regardless of whether a search finds the optimal graph or not, if a single graph is inferred and used for all downstream analysis, that point estimate would not intrinsically report confidence in various estimated features, such as the topology of relationships among populations, the presence or absence of admixture events, and the intensity of those events. There might be many graphs that fit the data equally well, and we should have more confidence in features shared among many of them than we should in features that are only found in some of them; shared features are most likely signals in the data while those that rarely occur are most likely spurious. Analyses based on a single graph do not distinguish between features that are estimated with high confidence and those estimated with low confidence. While it is possible to generate a distribution of TreeMix graphs across independent analyses of bootstrap replicates, it is rarely done in practice.

Here, we provide an alternative to greedy searches. Based on a model similar to TreeMix and qpGraph, we develop a Bayesian approach to sample over the graph-space using a reversible jump Markov Chain Monte Carlo (MCMC) method. The method can identify the graph with the highest likelihood encountered by the MCMC sampler, thereby effectively working as a heuristic maximum-likelihood optimizer, or it can report several summaries of the posterior distribution of admixture graphs. For example, it can estimate the graph topology with the highest marginal likelihood when integrated over admixture and divergence times as measured by occupancy in the MCMC sampler. Such a marginal likelihood is computed in *admixturegraph* ^13^ as well, but the exhaustive search algorithm of admixturegraph finds the graph with the highest likelihood - not the graph shape with the highest marginal likelihood. A particular strength of our new method is that it circumvents the need to report a single best graph by allowing calculations of posterior probabilities of particular marginal relationships between populations. We consider three approaches for this: one based on simplifying admixture graphs into simpler structures, one based on summarizing shared topologies into a consensus graph, and one based on subgraph analysis. If the number of leaves in the considered subgraph is kept small, we will observe few distinct subgraphs with these leaves, and we can estimate a complete posterior distribution over these graphs. Sampling subgraphs from the space of full graphs allows us to incorporate information from other populations when exploring the relationship between a subset of the populations.

We illustrate the utility of our method using simulations and reanalyze a previously published genomic dataset of Siberians and Native Americans^14^. We use the method to revisit two important and controversial questions in the history of the peopling of the Americas. First, we analyze the origin of the Inuit and show that they are modeled best as an admixture between a population represented by the Saqqaq genome, and Native Americans, represented by Athabascans. Secondly, we show that Athabascans are best represented as admixed between a Native American population and a Siberian population most closely related to the Koryak, but not the Saqqaq.

## Results

The Methods section describes our implementation of a Markov Chain Monte Carlo (MCMC) algorithm, *AdmixtureBayes*, which samples admixture graphs from their posterior distribution. We summarize genetic data from multiple populations as a matrix that captures how allele frequencies in the data covary between populations. AdmixtureBayes samples graphs that explain this covariance matrix. The topology of any sampled graph captures the relationships between samples as a mixture of the graphically structured covariance matrices. Branch lengths capture the amount of genetic divergence between populations, measured by drift, and admixture events explain shared allelic covariance between otherwise independently evolving populations. As a property of the MCMC algorithm, each graph is sampled at a frequency corresponding to its posterior probability. AdmixtureBayes is available to use at https://github.com/avaughn271/AdmixtureBayes.

### Evaluating convergence and mixing rate

In order to evaluate the convergence of the MCMC sampler, we used two different metrics, both of which are based on examining summary statistics of the chain. The summary statistics we chose to consider were the number of admixture events, the posterior probability, and the total branch length of the graph. Our first metric was simply examining the trace plots of the chain. From these plots, it is often possible to visualize the burn-in period. The second metric was the more sophisticated Gelman-Rubin convergence diagnostic, which analyzes the behavior of several chains run in parallel from different starting states.^15^ This diagnostic is based on calculating the ratio of the variance of the summary statistic between chains to the variance of the summary statistic within chains. A ratio close to 1 signifies that all chains have converged from their disparate starting states to the same equilibrium distribution. We used the *coda* package to perform this comparison.^16^ To evaluate the mixing rates of the chain, we plotted the autocorrelation of the summary statistics as a function of the lag between samples.

We demonstrated these analyses on a simulated dataset. Our simulated dataset was generated with msprime^17^ with a genomic region with mutation and recombination rates both equal to 10^−8^ and with 11 subpopulations. We used the following code snippet:

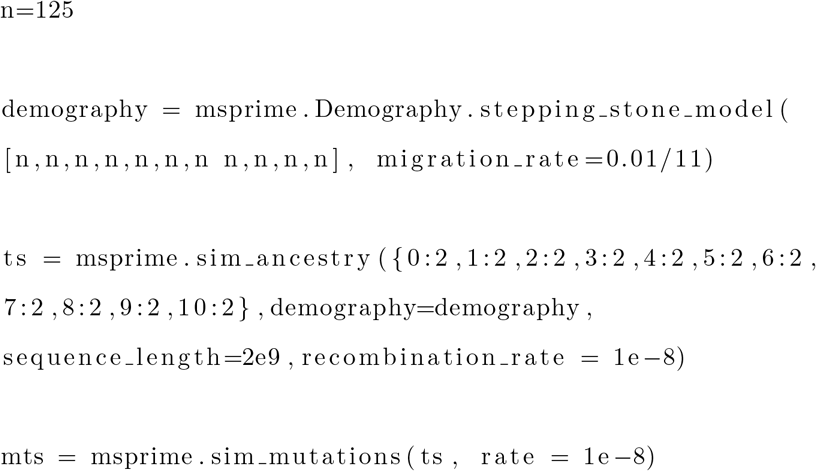

We took the first population to be the outgroup, which gives 10 leaf populations. We then ran AdmixtureBayes for three different chains, each with `--MCMC_chains 16` and `--n 150000` and using a random starting state, which is the default behavior of AdmixtureBayes. We plot the convergence and mixing results in Supplementary Figures S10, S11, and S12.

### Comparisons with TreeMix

To compare the accuracy of AdmixtureBayes to TreeMix, we first simulated several datasets using an admixture graph simulator (see Appendix A.1). The admixture graphs all had 10 populations and 0, 1, 2 or 5 admixture events. Based on the admixture graph, we simulated SNP data where we varied the number of haplotypes per population between 2, 10 and 50 using *ms* ^18^. Instead of simulating a fixed number of SNPs, we simulated datasets with a fixed mutation rate across 2000 or 5000 independent segments of 500 kb linked sites. This produced 250,000-850,000 SNPs, which we filtered to yield 25,000-85,000 effectively independent SNPs (see Methods), which is comparable in size to real biological datasets. Genomic datasets in humans, resulting from whole genome sequencing, typically contain information corresponding to between 50,000 and 100,000 effectively independent SNPs (see section on Saqqaq, Inuit, and Native Americans).

We then analyzed all simulated datasets with both AdmixtureBayes and TreeMix (see Appendix A.2). Comparing their accuracy is not straightforward because TreeMix produces one graph whereas AdmixtureBayes produces a posterior sample of graphs. In addition, TreeMix assumes a fixed number of admixture events, whereas AdmixtureBayes samples graphs with different numbers of admixture events. To solve the latter issue, we ran TreeMix conditioned on the true number of admixture events whereas we thinned the AdmixtureBayes samples such that they only contained admixture graphs with the true number of admixture events. However, to illustrate the ability of AdmixtureBayes to infer the number of admixture events, we include both the thinned and unthinned AdmixtureBayes samples in the plots. We used five metrics to compare the methods. The Mean Topology Equality is the proportion of the Markov chain spent in the true topology, which approximates the posterior probability of the true topology. The Mode Topology Equality is the proportion of replicates in which the maximum a posterior (MAP) estimate of the topology equals the true topology. The Mode Topology Equality and Mean Topology Equality were both compared to the proportion of times TreeMix infers the correct topology. The next metric we considered is the Covariance Distance, defined as the average of the Frobenius distance between the true graph and the covariance matrices implied by the MCMC-sampled graphs (see Methods). We compared that to the Frobenius distance between the covariance matrix implied by the TreeMix inferred graph and the true graph. Finally, we measured the Set Distance, which we defined as a topological distance measure similar to the Robinson-Foulds metric (Figure S8; Methods section). We evaluated both the ergodic average of the Set Distance between a graph in the chain and the true graph (Mean Set Distance) and the Set Distance between the MAP estimate of the topology and the true topology (Mode Set Distance). In both cases, we compared the results to the Set Distance between the true topology and the TreeMix inferred topology.

We note that the accuracy of the MAP estimate of the topology for a given number of admixture events is similar in both methods (Figure S1a), although AdmixtureBayes is perhaps slightly better when the number of admixture events is > 0. However, we also notice that neither method infers the true topology with high probability when the number of admixture events is equal to 2 or larger. This suggests that it may not be scientifically meaningful to focus on a single estimate of an admixture graph with 10 or more populations and 2 or more admixture events. Using Mode Set Distance, the story is somewhat similar (Figure S1b), but the advantage of AdmixtureBayes over TreeMix becomes more apparent in that the Mode Set Distance for AdmixtureBayes with 2 admixture events is considerably lower than the distance between the TreeMix graph and the true graph. In other words, when the methods infer incorrect topologies, the MAP estimates obtained by AdmixtureBayes tend in average to be less wrong than the estimates obtained by TreeMix. Because both methods use similar evolutionary models, improved optimization in TreeMix would likely lead to a performance more similar to that observed for AdmixtureBayes. The three metrics evaluating ergodic averages over the chain tell slightly different stories (Figure S1c-e). While the Mean Set Distance is still considerably smaller for AdmixtureBayes than for TreeMix (Figure S1d), the ergodic average of the Topology Equality is slightly smaller (Figure S1c), and the Covariance Distance is slightly larger (Figure S1e) for AdmixtureBayes. That is, while a randomly sampled graph from the posterior tends to be closer (in terms of Set Distance) to the true graph than the TreeMix estimate is, the same is not true when using Covariance Distance or Topology Equality. It is expected that the MAP estimate is more accurate than the average posterior graph, yet a large difference could be a sign that the sampled posterior distribution is inaccurate. However, the difference here is small, which supports the correctness of the posterior sampling.

We varied the sample sizes (number of individuals per population used to estimate allele frequencies), to determine the dependence of these conclusions on sample size (see Figure S2). In general, the conclusions seem to follow those obtained in the previous simulations. It is also apparent that there is a pronounced advantage in terms of accuracy in moving from 2 to 10 haplotypes per population. The improvement in performance is smaller when moving from 10 to 50 haplotypes.

### Summarizing subgraphs

While it may be difficult to obtain a unique point estimate of the admixture graph with highest statistical support, particularly for analyses involving many populations and large state spaces of graphs, elements of the graph may, nonetheless, be well supported. It is possible to consider the relative support, in terms of posterior probability, of individual subgraphs. Analyzing the support for subgraphs within the context of a larger admixture graph has an advantage over analyses limited to the focal populations represented in the subgraph, that information from other populations can be directly taken into account.

Using parts of the same simulations as before (see Appendix A.1), we explored the accuracy of subgraph inference. For each dataset, we considered subgraphs for 3, 4 and 5 randomly drawn focal populations. Having already analyzed the datasets for all 10 populations, we extracted estimates by marginalizing the estimates for the full admixture graph, which we denote as subgraphs from the ‘Big’ dataset. We also recomputed the subgraphs by analyzing only the data from the focal population while discarding all information from the non-focal populations, which we denote as ‘Small’ graphs. Marginalizing a joint graph is presumably better than estimating the marginal graph, because the joint estimation uses information from all populations of the dataset. As expected, the Big graphs estimated by AdmixtureBayes do have higher accuracy than the Small graphs (Figure S3). Surprisingly, the same is not the case for TreeMix Big and Small graphs. This pattern is repeated for most accuracy and distance measures and subgraph sizes (Supplementary Figure S14). TreeMix graphs are more accurate for very small graphs, but they lose accuracy fast as the number of populations increases (Figure S14, Small columns). Figure S3 suggests that the expected improvement of the Big TreeMix subgraphs compared to the Small subgraphs decreases when including more populations.

### Exploring the genetic history of Saqqaq, Inuit and Native Americans

We applied AdmixtureBayes to a set of previously published Siberian and Native American samples^14^ to explore the relationship between Siberian Chukotko-Kamchatcan speakers (Koryak), an ancient individual from the extinct Saqqaq culture (Saqqaq), Inuit-Yupik-Unangan speakers (Greenlandic Inuit), and Na-Dene speakers (Athabascan). The dataset also contained North and South Americans (Anzick, Aymara) and various other groups. We chose the Yoruba population as the outgroup. Running time of AdmixtureBayes was 50 hours in parallel on 32 cores.

To extract information from the posterior distribution of admixture graph topologies, we introduce two ways of summarizing relationships among sets of focal populations (for details, see Methods). Both are based on summarizing each sampled admixture graph in the posterior into a *topology set*, which is the set of all nodes labeled by their descendants. This discards information about the number of and timing of admixture events (see Figure S8). From such a topology set, we can create the *minimal topology*, which is the ‘simplest’ directed graph yielding the same topology set (see Figure S9). The two minimal topologies with the highest posterior probabilities are shown in Figure S4. We also considered the frequency of each internal node across posterior samples. In Figure S4 these frequencies are denoted as percentages in parentheses in each node. The second summary of the admixture graph sample is the set of nodes with a frequency higher than *α* in the topology sets, which we denote as the *consensus graph* at threshold *α*. Figure S5 shows this summary for *α* = 0.75.

While no single graph received high support when including all data, we can extract subgraphs that are informative about the relationships between specific subsets of populations. In particular, there has been considerable debate about the relationships between populations represented by the Koryak, Saqqaq, Greenlanders, and the Athabascans. Archaeological evidence suggests that the Inuit people from Greenland and people from the now extinct Saqqaq culture represent independent migrations into the Americas from Eastern Siberia and the area around the Bering strait ^19 20 21^. However, there is some debate about the origin of the Athabascans^22 23 20 24^. Most molecular evidence of Athabascan ancestry is thought to have originated from the first migration of people into the Americas that also gave rise to most other Native American groups, such as the indigenous people in Central and South America. However, some portion of genetic variation in Athabascans seems to have also originated from other groups, perhaps related to Inuit, Saqqaq, or other Siberians such as the Koryak. Naming and identifying sources of genetic variation is further complicated by the fact that these possible reference populations may themselves be admixed. A marginal analysis of the relationship between Koryak, Saqqaq, Greenlanders, and Athabascans, that can take gene flow from other groups into account, is therefore very much wanted.

Figures S6 and S7 depict the subgraphs for different subsets of these groups and for all groups together, extracted from the posterior distribution of graphs from the full dataset. The most strongly supported subgraph for Saqqaq, Athabascan, and Koryak supports the tree ((Athabascan, Koryak), Saqqaq) with 96% posterior probability. This implies that a relationship where the gene flow into Athabascans came from a population closer to the Saqqaq, than to the Koryak from Siberia, is not supported by the data. In contrast, when considering the relationship between Koryak, Athabascans and the Inuit Greenlanders, the most strongly supported admixture graph is a tree with the structure ((Athabascan, Greenlander), Koryak), likely reflecting gene flow into the Inuit Greenlander from Native Americans related to Athabascans. We emphasize that in these inferences, by analyzing the posterior probability of subgraphs embedded within larger graphs, we have also explicitly modeled the effects of gene flow from other groups including various Siberian, Native American, and East Asian groups. When considering all four populations together, the Greenlanders are best modeled as a population admixed between Athabascan related populations and Saqqaq related populations. Again, there is no apparent gene flow between the Saqqaq and the Athabascans following their initial divergence.

## Discussion

We here present the program AdmixtureBayes, which is a method for inferring admixture graphs using MCMC. On simulated data, it infers graphs more accurately than TreeMix under the Set Distance measure. AdmixtureBayes also tends to find the true topology with slightly higher probability than TreeMix does. We speculate that the superior performance of AdmixtureBayes in terms of Set Distance is likely caused by issues relating to optimization in TreeMix. Possibly, the procedure in TreeMix could be improved to more or less match the performance of AdmixtureBayes in terms of producing point estimates. However, we also note that for larger graphs, the probability that the graph identified is the true graph is very small for both methods. This suggests that the reporting of a single graph may not necessarily be accurate. As is common practice in phylogenetics, admixture graphs should report measures of statistical confidence for the relationships inferred among internal nodes in the graph, as is reported in this paper. We also encourage the use of embedded subgraphs as a powerful approach for investigating the relationship between specific populations while taking gene flow from other reference populations into account. The use of posterior probabilities, as reported here, is facilitated by the use of a bootstrap procedure that can estimate the effective number of independent SNPs. In our real data analysis, we obtained information from human genomes corresponding to approximately 40,000 independent SNPs. This number determines the peakedness of the likelihood surface, which directly influences the posterior distribution of admixture graphs. TreeMix and qpGraph employ similar resampling techniques to obtain variance estimates that control the peakedness of their likelihood surfaces, thereby reducing the complexity of admixture graphs explored during inference.

Our analysis of Native American and Siberian samples largely recapitulates many previous analyses and identifies many admixture events ^14^. Furthermore, we find a similar, but not identical topology, to a previous admixture topology^14^. However, our results also indicate that several features of the true admixture graph remain uncertain. For example, we could not definitively resolve the question of introgression into the Han lineage from the ancestral lineage of Ust’-Ishim. Our analysis does not support previous claims that the Saqqaq culture is a good proxy for the source of gene flow into Athabascans^23 20^, although statistical power could still be improved. In both our analysis and previous work, each population is represented by just one or two diploid individuals. Our simulations suggest that increasing the number of individuals per population might lead to substantially improved statistical accuracy. We also note that the sample quality was relatively poor for some samples analyzed here, particularly the Saqqaq, which has many missing sites.

The estimation of admixture graphs is becoming one of the most important tools in population genomics. However, methods for estimating such graphs are still in their infancy. AdmixtureBayes provides a step towards improved estimation and more rigorous quantification of statistical uncertainty in admixture graph inference.

## Methods

### Data

We analyzed a dataset consisting of SNPs for 12 human populations that was first analyzed by Moreno-Mayer et al.^14^. We treated the Yoruba population as an outgroup leaving effectively 11 populations with unknown relationships to estimate. One diploid individual was sampled from each population, except the Koryak, Ket, Greenlander and Athabascan populations, which each had two diploid individuals. Whole genome-sequencing was performed on each individual to provide an average coverage between 1X (for the Malta individual) to 44.2X (for one of the Greenlander individuals). Further details regarding sequencing and data processing methods are described in Moreno-Mayer et al.^14^. The alleles for the ancient individuals from the populations Saqqaq, Malta, Anzick and USR1 that were not transversions were treated as missing. We then filtered out any site for which there was a population with missing data. In total 251,542 biallelic SNPs were retained. Large numbers of missing SNPs for some individuals is not a computational problem for AdmixtureBayes, though it does violate the assumption of even sampling imposed by the Wishart distribution (see Methods, equation (5)).

### AdmixtureBayes Model

The AdmixtureBayes program searches the posterior distribution of admixture graphs given observed SNP data using a Markov Chain Monte Carlo procedure. To assess the likelihood of an admixture graph we summarize both the admixture graph and the data as covariance matrices of allele frequency changes^2^. The admixture graph covariance matrix is calculated as in TreeMix. Consider the tree structure in Figure 1 where population 2 is a mix of two ancestral populations with proportions *w* and 1 − *w*.

**Fig 1:**
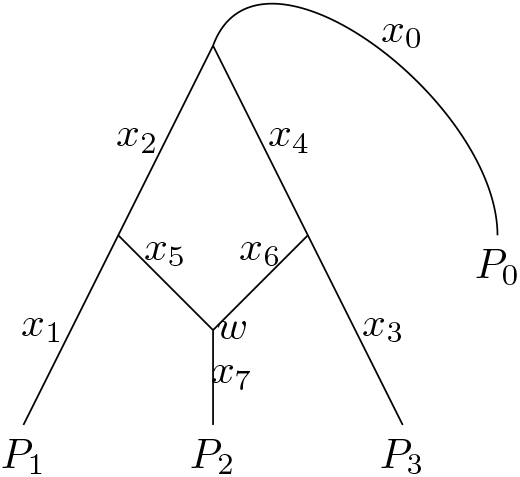
An admixture graph for the 3 populations and one outgroup. Considering a single SNP, the quantities *x*_1_, …, *x*_7_ are changes in allele frequency, *w* is the admixture proportion, and *P*_0_, *P*_1_, *P*_2_ and *P*_3_ are allele frequencies in the sampled populations.

The allele frequency in the 4 populations, *P*_0_, *P*_1_, *P*_2_ and *P*_3_ are related through the allele frequency changes *x*_0_, …, *x*_7_ at any SNP.

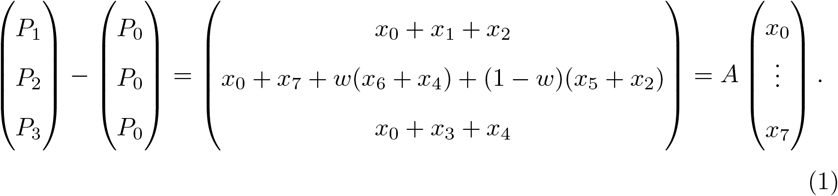

Notice that *A* is a matrix that only depends on the admixture graph through the graph structure and admixture proportions. We consider the vector of allele frequency drifts terms (*x*_0_ … *x*_7_) to be stochastic because it depends on a random sample of SNPs. In the neutral Wright-Fisher model, changes in allele frequencies due to genetic drift can be approximated by a normal distribution when the allele frequency change is small and the frequency is far from the boundaries at 0 and 1. If *x*_*i*_ is the amount of drift from a node with allele frequency *p*_*i*_, then the allele frequency change can be approximated as 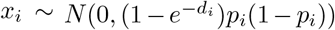 where *d*_*i*_ = *t*_*i*_*/*2*N*_*i*_ is the number of generations scaled with the population size ^25^. We collect the factor 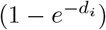 into a single factor *c*_*i*_ and substitute the node-specific *p*_*i*_ with a SNP-global *p* giving the tractable, approximate, expression

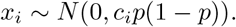

Consequently, we can approximate the joint distribution of allele frequencies at all leaf nodes as

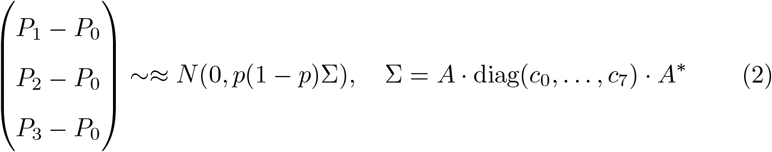

where matrix Σ is called the *admixture graph covariance matrix*.

The empirical estimate of the covariance of allele frequencies is denoted the *data covariance matrix*. In real data we never observe the population allele frequencies but rather the sample allele frequencies. This complicates the computation of the data covariance matrix slightly. Let *p*_*ij*_ be the sample allele frequency in the *i*’th population at the *j*’th SNP, *i* = 0, 1, …, *n, j* = 1, …, *N*. They are assumed to come from the distribution

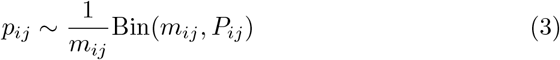

where *m*_*ij*_ is the number of haplotypes sampled and *P*_*ij*_ is the population allele frequency.

Denote population *i* = 0 an outgroup, and consider the intuitive estimate of the covariance matrix

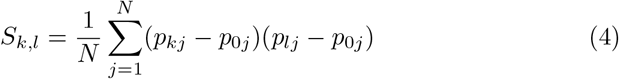

If there are any missing values in a summand, we leave that summand out of the sum. Regardless of missing values, (4) is inherently biased because the inner term (*p*_*kj*_−*p*_0*j*_)(*p*_*lj*_−*p*_0*j*_) does not have the same mean as (*P*_*kj*_−*P*_0*j*_)(*P*_*lj*_−*P*_0*j*_). From (3) we calculate the difference as

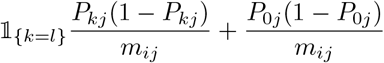

which suggests the following bias correction factor for *S*_*k,l*_:

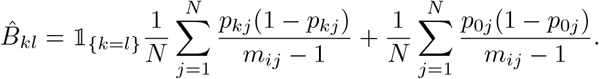

After correcting, we normalize with

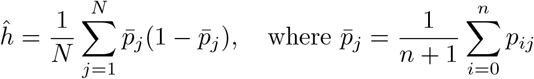

to take the factor *p*(1 − *p*) from (2) into account.

If the sample allele frequencies were normally distributed and independent across markers, the estimator in (4) would be Wishart distributed and the degrees of freedom would be the number of markers. The sample allele frequencies are not independent and only approximately normal, yet we use the likelihood

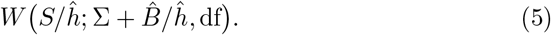

The degrees of freedom, df, is adjusted to take into account the lack of independence. We estimate df using *R* bootstrapped replicates of 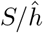 which we will denote *X*^(1)^, …, *X*^(*R*)^. Let 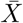 be the average of the bootstrap samples. It would be natural to estimate the df with the maximum likelihood of the model

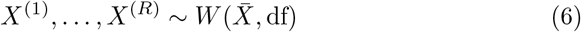

However, simulations show that the estimates of df from (6) give results that are less accurate than the following moment-based estimator (Supplementary Figure S13). We take advantage of the fact that the variance of the (*k, l*)’th entry of a Wishart distribution with mean Ψ/df and degrees of freedom, df, is

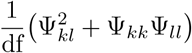

to estimate the df as

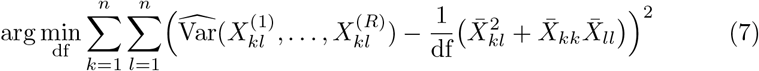

where 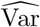 is the sample variance. This moment-based estimator leads to better performance of AdmixtureBayes (Supplementary Figure S13).

In practice, to make the inference more robust to deviations from the prior, we normalize the matrices by using the likelihood

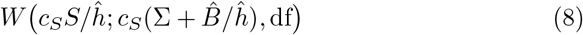

where 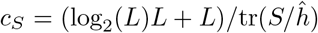. For more on this, see the later section on Robustness Correction.

### Admixture Graphs

An admixture graph consists of a topology and a set of continuous parameters. The space of topologies for a given number of leaves, *L*, consists of all uniquely labeled graphs of the set of all directed acyclic graphs which fulfills

1. There exists one and only one root. That is a node with no parents and exactly two children.
2. The number of nodes with no children is *L*. All these nodes have only one parent and are called leaves.
3. If a node is not a root nor a leaf, it has either
  a. 1 parent and 2 children in which case we call it divergence node.
  b. 2 parents and 1 child in which case we call it admixture node.
4. There are no *eyes*, i.e. the parent nodes of an admixture node are distinct (and the child nodes of a divergence node are distinct).

The labeling consists of

1. All leaves are given a unique label.
2. Parent edges of an admixture node can be either a ‘main’ branch or an ‘admixture’ branch. All admixture nodes have one parent edge of each type.

We do not label branches and nodes in general, meaning that even though the the leaves are given a unique label, the leaves themselves are not unique. For example, switching the labels of two leaves that form a cherry in the graph, would not change the graph. For a more formal definition, see the definition of topology in Appendix A.3.

All branches have a length in the interval (0, ∞) and all admixture nodes are given an admixture proportion in the interval (0, 1).

### Prior

We define a prior on the topology, 𝒢, and on the continuous parameters of the admixture graph. The continuous parameters include the branch lengths, 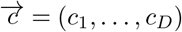, and the admixture proportions 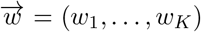. Let *K* denote the number of admixture events, *L* the number of leaves, and *D* = 2*L* − 2 + 3*K* the number of branches. The full prior is then

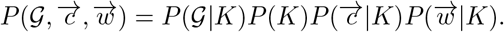

The prior on the number of admixture events is a geometric distribution with parameter 0.5 (truncated to max 20). The prior on 𝒢, *P*(𝒢|*K*), is a uniform prior on all labeled admixture graphs with *K* admixture events. To evaluate this prior, we need to calculate the number of possible topologies for a given number of admixture events. Therefore we have derived the recurrence formula

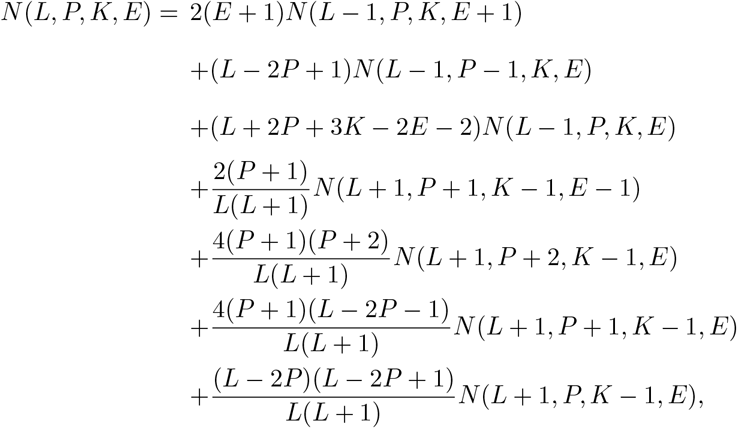

where *L* is the number of leaves, *P* is the number of pairs of leaves that share a common parent, *K* is the number of admixture events, *E* is the number of eyes, and *N*(*L, P, K, E*) is the number of unique topologies with those attributes. Notice that we here allow eyes which otherwise are disallowed in our definition of admixture graphs. See Appendix A.3 for proof. Then

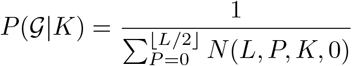

For the admixture proportion prior, 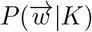, we chose the uniform distribution on the interval (0, 1).

For the prior on the branch lengths, 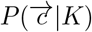, we chose to let all branch lengths be independent and marginally follow the distribution

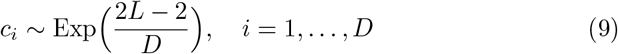

The rate of the exponential prior adapts to the topology such that graphs with many branches, and thereby many admixture events, are expected to have smaller branch lengths. For motivation see the following section on Robustness Correction.

### Robustness Correction

In the Bayesian phylogeny program MrBayes^26^, it has been shown that independent, exponentially distributed priors on the branch lengths can unduly influence posterior estimates of total tree length^27^, which could also be a problem for AdmixtureBayes. To see this, consider the average branch length 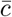. For simplicity, assume the effective population size, *N*_*e*_, is constant across the admixture graph. Furthermore, suppose that the exponential rate of (9) is 1. Let *T* = ∑*T*_*i*_ be the total *time*(not drift) of all branches in the admixture graph. Then we can write

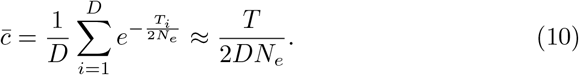

Since it is an average of independent random variables, its mean and variance are

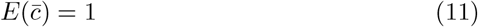

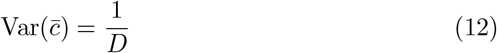

This means that the prior expects 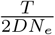 to be very close to 1. However, for real datasets we would expect the ratio to vary much more, and there is no biological reason why it should be near the arbitrary number 1. For a specific dataset, if the true value of 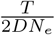 were smaller than 1, the posterior would be overestimated for admixture graphs with higher values of 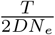. Such graphs would generally possess a deflated number of admixture events and thereby a smaller *D*. Similarly, large true values of the ratio would result in a skew towards admixture graphs with an inflated number of admixture events.

To mitigate the problems caused by the independent, exponential priors, Mr-Bayes includes an alternative compound Dirichlet-Gamma prior on the branch lengths, such that the variance of the average branch length can be set arbitrarily high^27^. However, we normalize the data covariance matrix and adjust the rates of the exponential distributions accordingly.

To reduce the sensitivity of our posterior estimates to the prior, we wish for the prior exponential rate of *c*_*i*_ to be close to 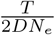. We rewrite

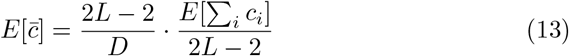

The first fraction is manageable because the prior is allowed to depend on *L* and *D*. The second fraction is the average branch length if there are no admixture events in the admixture graph. It can be estimated by summing the outgroup-leaf distances for all leaves and dividing by the number of branches between the outgroup and the leaves. Denote that divisor 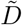. Unfortunately, 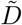 depends on the topology. Therefore, we approximate 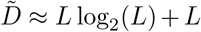, which leads to the approximation

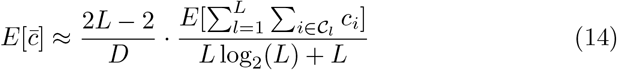

where 𝒞_*l*_ is the set of indices of the branches between the outgroup and leaf *l*. Regardless of the true topology, we can estimate 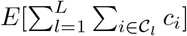 by the trace of the data covariance matrix.

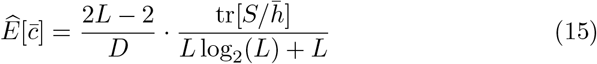

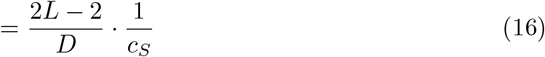

Instead of letting (16) be the exponential rate of the branch length prior, we normalize the data covariance matrix by *c*_*S*_ and let 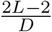 be the expected mean of the branch lengths. We avoid having a prior that depends on the data by moving *c*_*S*_ out of the prior. However, since *c*_*S*_ depends on the data, the matrix 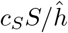 would not be Wishart distributed, even if 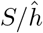 were truly Wishart distributed. The scaling by *c*_*S*_ therefore adds another layer of approximation to the likelihood.

This robustness correction makes the graph inference independent of the absolute scale (as measured by the trace) of the data covariance matrix. The maximum likelihood methods TreeMix^2^, qpGraph^1^, and MixMapper^3^ inherently have this property as well.

### MCMC

The MCMC is implemented as a parallel Metropolis coupled MCMC algorithm^28 29^ to increase the number of jumps between modes of the posterior surface. Because admixture graphs with different number of admixture events also have different numbers of continuous parameters, we use the reversible jump generalization of the MCMC algorithm^30^. The proposal distribution is a mix of 7 smaller proposals. They are

1. Add an admixture branch to the admixture graph. An admixture branch goes from a *source branch* to a *sink branch*(Figure 2). To make the proposal, a random sink branch, *s*, is chosen with probability 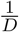 where *D* is the number of branches in the graph (not including the branch to the outgroup). Next, a random source branch, *s*′, is chosen from the remaining branches (including the root/outgroup branch) such that an addition of an admixture branch would not create a cycle in the graph. If the number of possible sink branches is *D*′(*s*), the probability of the sink position is 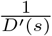. Next the attachment point on the sink branch is simulated uniformly. If the branch lengths of *s* and *s*′ is *c*(*s*) and *c*(*s*′) the attachment outcome has density 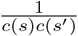. If the source branch is the root branch, we simulate the attachment point with an exponential distribution, Exp(1), instead. The new admixture proportion is simulated uniformly between 0 and 1, and the admixture branch length, 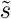, is simulated from Exp(1) with density 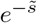. Lastly, the labeling of the two parent branches of the new admixture node is simulated. The probability of either possible labeling is 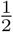. In conclusion, the density is

**Fig 2:**
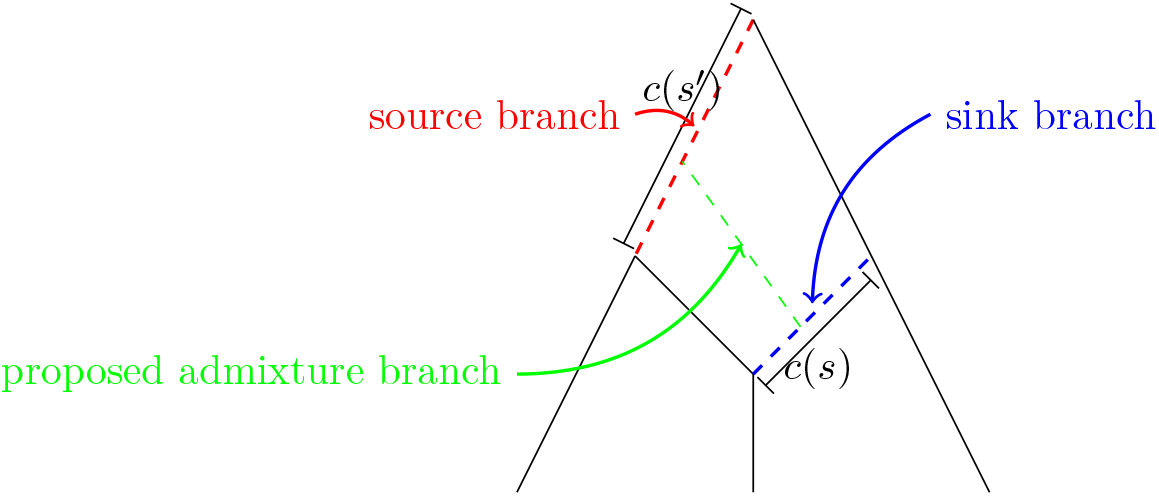
When adding an admixture branch (green), we will randomly draw the branch where it comes from, the source branch (red). The admixture branch goes into the sink branch (blue).

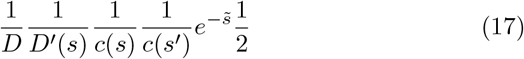 To find the acceptance probability of this proposal, we calculate the proposal probability of the reverse move (see proposal number 2). The reversible jump Jacobian factor is 1.
2. Remove an admixture branch from the admixture graph. An admixture branch can be removed if 1) its parent is not an admixture node and 2) its removal will not cause an eye. Let the number of admixture branches eligible for removal be *K*′. We choose uniformly from that set and remove the admixture branch. The density is

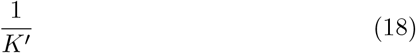
3. Node sliding. A random branch whose parent is a divergence node is chosen. We move its attachment point to its source branch a distance λ*x* where *x* ∼ *χ*^2^(1). A node can often be slid either up and down and sometimes the sliding node meets a bifurcation where it can slide in either of two directions. We choose the new node position uniformly from the set of the possible sliding destinations, following the topological constraints defined in step 1. If the sliding node slides out of the graph, we reject the proposal. The forward density is 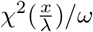 where *ω* is the number of possible sliding destinations for a node when moved a distance *x* from its position in the current graph. We compute the backward density using the same formula. We update λ on-the-fly following guidelines for adaptive proposals in MCMC^31^, eliminating the need for pre-analysis parameter tuning.
4. Random walk on the branch lengths. We add a normally distributed noise to all the branch lengths. If any branch length become negative, we automatically reject the proposal. The backward density is identical to the forward density. The variance of the random walk increments is controlled by parameter *s* which we also adapt on-the-fly using adaptive strategies.
5. Random walk on the admixture proportions as in step 4 but with another *s*-value. Proposals outside (0, 1) are rejected.
6. Random walk on the branch to the outgroup as in step 4 but with another *s*-value. Negative proposed branch lengths are again rejected.
7. Random walk on the branch lengths but inside the null space of matrix *A*. This means that the proposed admixture graph will have the same covariance matrix - and therefore the same likelihood - as the previous graph. This proposal is also adaptive, as in step 4.

### Graph Summaries

In the Results section we explained the two summaries, *minimal topology* and *consensus graph*, which we will define formally here. Furthermore, we introduced the Set Distance used to measure distances between admixture graph topologies and the Covariance Distance for distances between admixture graphs. In this section, we define these quantities.

The Covariance Distance between two admixture graphs with *L* leaves and covariance matrices Σ^1^ and Σ^2^, respectively, is

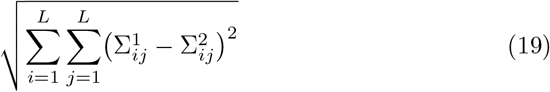

For a single node let the *descendant set* be the the set of its leaf descendants, e.g. *t* = {*l*_1_, *l*_2_, …, *l*_*a*_}. For a topology, let *T* be the *topology set*, which is the set of descendant sets of all its nodes, excluding those sets for the leaf and root nodes. The *minimal topology* is the extension of such a topology set to a directed graph. The extension starts by adding the trivial descendant sets for the leaves (containing only one leaf) and the root (containing all the leaves). Denote this set 𝒯. The minimal topology has the same nodes as 𝒯 and there is a connection from node *t* ∈ 𝒯 to *t′* ∈ 𝒯 if

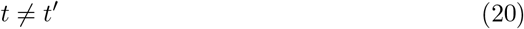

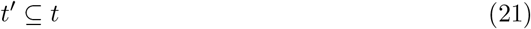

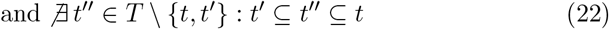

To summarize a sample of admixture graphs, *g*_1_, …, *g*_*R*_, using a consensus graph, we first transform all of them into their topology sets and obtain a sample *T*_1_, …, *T*_*R*_. The posterior probability of a node can be estimated by the sample frequency

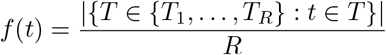

The topology set of the *consensus graph* at threshold *α* is

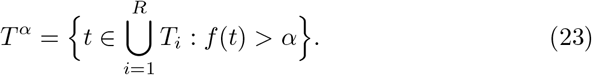

The consensus graph itself is obtained by extending *T*^*α*^ to a directed graph with the rules (20)-(22).

The *Set Distance* between two graphs *g*_1_ and *g*_2_ with topology sets *T*_1_ and *T*_2_ is

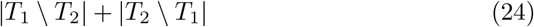

## 0.1 Author Contributions

T.M and R.N conceived the idea and initiated the project. The mathematical framework was developed by R.N, S.N and M.L. K.L derived and proved the formula for the number of topologies. S.N and A.V implemented code and simulations. Planning and interpretation of simulations and data analysis was done by S.N, A.V, R.N and T.M. S.N, A.V, and R.N wrote the manuscript with input from all authors.

## A Appendix

### A.1 Simulations of admixture graphs and datasets

In the simulation studies in Figures S1, S2, S3, S14 and S15, we have simulated admixture graphs. Their number of admixture events, admixture proportions and branch lengths are simulated from our prior. However, our uniform prior on the topologies does not naturally yield a simulation method. We constructed an alternative algorithm that simulates admixture graphs conditioned on the number of admixture events using a discrete-time Markov chain that follows lineages back in time. If there are *L* leaves, there are *L* free lineages at the start. Given the number of leaves and the number of admixture events, we know the number of divergence and admixture nodes. The free lineages choose a parent node uniformly at random such that

1. No more than two lineages choose the same divergence node
2. No more than one lineage choose the same admixture node
3. No ‘eyes’ are formed. That is, two lineages from the same admixture node will not choose the same divergence node.
4. The complete admixture graph can still be constructed. For example, if no two lineages had chosen the same divergence node, there would not be any free lineages in the next step of the Markov chain.

When two lineages have chosen a divergence node, a new free lineage is released for the next Markov chain. Likewise, a chosen admixture node produces two new lineages. The algorithm stops when there is just one free lineage left and all divergence nodes and admixture nodes have been ‘filled’. For topologies without admixture events, our simulation algorithm chooses uniformly between the possible topologies. For topologies with admixture events, the algorithm prefers admixture events closer to the root when compared to the uniform prior.

TreeMix and AdmixtureBayes do not operate with the same admixture graph space. TreeMix allows admixture flow into the outgroup population, whereas AdmixtureBayes does not. On the other hand, AdmixtureBayes searches through admixture graphs with *invisible admixture events*. An admixture graph has an invisible admixture event if its admixture graph covariance matrix can be obtained by another admixture graph with fewer admixture events. For a fair comparison, all simulated admixture graphs are from the intersection of the two admixture graph spaces.

The computations behind Figures S1, S2, S3, S14 and S15 also include simulation of genetic data using *ms* ^18^. A command given to *ms* for simulation of a graph with 5 populations could be

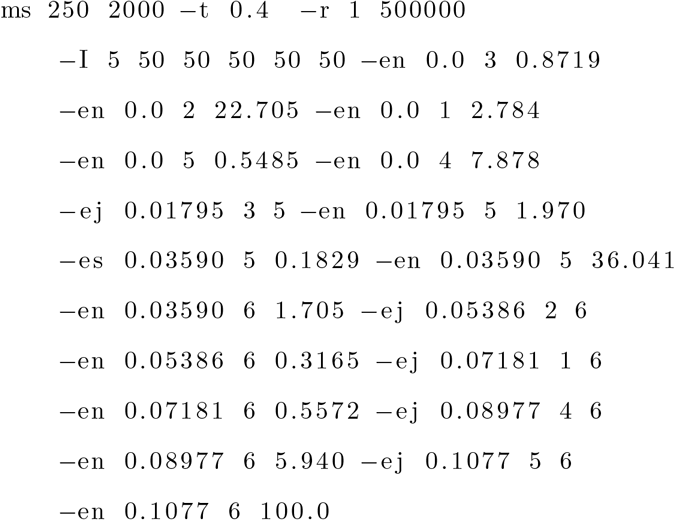

In the above code 50 individuals are sampled in each population. To assess how well AdmixtureBayes and TreeMix handle linked sites, we always simulate linked SNPs (with *r/t* = 1*/*0.4 = 2.5). To lessen the computational burden, we simulate the genome in 2000 independent segments of 500,000 sites. The branch lengths of the admixture graph are incorporated by adjusting the population sizes using the “-en” option. We always set the root population size to the high value 100 to increase the number SNPs present in all populations. The outputs of our *ms* commands are easily transformed into the input format of TreeMix (which is identical to the input of AdmixtureBayes).

In the simulation study for comparing AdmixtureBayes and TreeMix on admixture graphs with 10 populations, we simulated independent datasets. All datasets were based on different admixture graphs with either 0, 1, 2 or 5 admixture events. The datasets also varied in the number of haplotypes (2, 20 or 50) and the number of genomic segments (2000 or 5000). We analyzed all datasets with both AdmixtureBayes and TreeMix.

In the simulation study for comparing AdmixtureBayes and TreeMix on subsets of admixture graphs with 10 populations, we simulated datasets with 0, 1 and 2 admixture events, 5000 independent DNA segments, and 10 haplotypes per population. For subsets of size *x*, we chose *x* populations uniformly from the 10 possible populations. From the AdmixtureBayes samples in the previous simulation study we extracted the marginal distributions of the subgraphs of those *x* populations. We compared those to the true subgraphs using Set Distance and Topology Equality. From the TreeMix estimates of the previous section we extracted subgraphs of the maximum likelihood admixture graphs and also compared them to the true subgraphs. We refer to both these types of results as ‘Big’ because they are based on the full, larger dataset. Likewise we extracted datasets for the same *x* populations and analyzed them with both TreeMix and AdmixtureBayes. We varied *x* between 3, 4 and 5 populations. A subgraph of an admixture graph will often contain a smaller number of admixture events than the full graph. Furthermore, the remaining admixture events may be invisible in the subgraph. TreeMix does not consider graphs with invisible admixture events. Therefore, we ran TreeMix with the number of visible admixture events in the subgraph. When the true subgraph contains invisible admixtures, TreeMix has probability 0 of finding the true graph. In comparison, AdmixtureBayes can rarely visit graphs with invisible admixture events, which could give it an unfair advantage in the comparison. Fortunately, this only influences the Mean Topology Equality and Mode Topology Equality of the Small subgraphs (not shown in Figure S3 but in Figure S14), because the Set Distance measures are unaffected by invisible admixture events.

### A.2 Running TreeMix and AdmixtureBayes

TreeMix can estimate a maximum likelihood graph for a fixed number of admixture events, but the higher the number of admixture events, the higher the maximum likelihood value. Therefore, the original TreeMix paper suggests iteratively adding admixture events and stopping when the added admixture event does not pass a test for statistical significance. However, to simplify the comparison, we ran TreeMix with the true number of admixture events. Otherwise we used the default settings of version 1.13. The program first estimates an initial admixture-free tree by iteratively adding best fitting populations in a random procedure. Next, the admixture branches are added deterministically. Because of the randomness of the first step, the starting seed could influence the results. However, preliminary results showed that repeating the TreeMix maximum likelihood optimization for different seeds and choosing the highest likelihood graph amongst the repeated analyses did not change the accuracy of the estimated admixture graphs when analyzing our simulated datasets. Most seeds produced the same maximum likelihood graphs. Therefore, we used only one seed.

For each analysis, AdmixtureBayes was run for up to 12 hours on 15 cores. We ended the analysis when the effective sample sizes of several summary statistics exceeded the threshold 200^32^ after removing the first half of the samples as burn-in. This is an indication that the Monte Carlo Markov Chain has adequately approximated its stationary distribution, which is the target posterior density. For this reason, many analyses lasted no longer than 1 hour. Even if a chain did not fulfill the stopping criteria after 12 hours, we included the chain in the subsequent analysis after removing the burn-in period. Otherwise, we used the default settings.

### A.3 Number of admixture graph topologies

In order to compute the prior on the space of admixture graphs, we use the number of possible admixture graph topologies with *K* admixture events. This number grows at least exponentially with *K* and is further complicated by our specific requirements to the admixture graph topology. For computational convenience we will consider an extended class of admixture topologies: a *multi-graph topology* with *L* leaves is an acyclic directed multigraph (which is a graph that allows more than one edge between two vertices) for which

1. There exists one and only one root. That is a node with no parents and exactly one child.
2. The number of nodes with no children is *L*. All these nodes have only one parent and are called leaves.
3. If a node is neither a root nor a leaf, it has either

a. 1 parent and 2 children in which case we call it a *divergence node*, or
b. 2 parents and 1 child in which case we call it an *admixture node*.

This extends our original definition of an admixture graph topology by allowing eyes, i.e. admixture nodes whose parent branches merge in the same divergence node. The root is also now a node with one child instead of two, which means that all multigraph topologies have a single branch “on top.” Furthermore, we explicitly label all inner nodes. As before each admixture node will have one *main* parent branch and one *admixture* parent branch. We will use the notation

- The edges leading to leaves are referred to as *terminal edges*.
- A set of two terminal edges from a single node is a *pair*.

These graph elements are illustrated in Figure 3.

**Fig 3:**
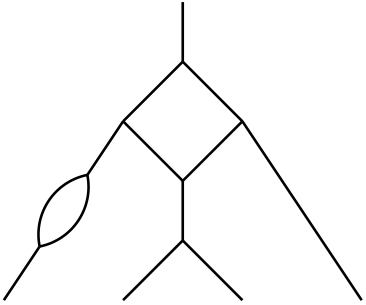
In all of our illustrations the direction of edges is from top to bottom, unless marked otherwise. This multigraph topology has 4 leaves, 1 pair, 2 admixture nodes, 5 divergence nodes, and 1 eye. The root is the node at the very top of the topology. We have not explicitly labeled the nodes and branches.

A multigraph topology consists of a set of nodes 𝒱, a set of main edges *ε*_*M*_, and a set of admixture edges *ε*_*A*_. There are *L* leaf nodes, {*l*_1_, …, *l*_*L*_} ⊆ 𝒱. For every admixture node, one of its parent branches belongs to *ε*_*A*_ and the other belongs to *ε*_*M*_. Note that all nodes are uniquely labeled. However, we are only interested in counting the number of topologies that differ in a nontrivial way. For example, switching the labels of leaf nodes that form a pair can be considered a trivial change to a topology. Therefore, we construct equivalence classes on the set of multigraph topologies and count those equivalence classes instead.

Let *ε* = *ε*_*A*_ ⊎ *ε*_*M*_ be the multiset union of *ε*_*M*_ and *ε*_*A*_. The admixture edges of a multigraph topology (𝒱, *ε*_*M*_, *ε*_*A*_) are classified into two subsets, *ε*_*M*_ and *ε*_*A*_, but we can also disregard the classification and consider the *reduced multi-graph topology*(𝒱, *ε*). We call a graph isomorphism between reduced multi-graph topologies *shape preserving* while a graph isomorphism between multi-graph topologies is *symmetry preserving*. A symmetry preserving isomorphism is clearly also shape preserving. If *f* is a symmetry preserving graph isomorphism, we say that *f* is *leaf preserving* if *f*(*l*_*j*_) = *l*_*j*_ for all *j* = 1, …, *L*. When counting admixture graphs, we consider two admixture graphs different if and only if they are not isomorphic under such an isomorphism.

For a fixed number of leaves *L*, number of pairs *P*, number of admixture events *K*, and eyes *E*, we will consider the three sets

1. The set of equivalence classes under shape preserving isomorphisms is denoted 𝒮_*L,P,K,E*_. The equivalence classes are called *shapes*.

2. The set of equivalence classes under symmetry preserving isomorphisms is denoted 𝒰_*L,P,K,E*_. The equivalence classes are called *unlabeled topologies*.

3. The set of equivalence classes under leaf preserving isomorphisms is denoted 𝒯_*L,P,K,E*_. The equivalence classes are called *topologies* and sometimes explicitly *labeled topologies*.

We are particularly interested in the cardinality of the set 𝒯_*L,P,K,E*_, which we denote by *N*(*L, P, K, E*). The difference between the sets 𝒮_*L,P,K,E*_, 𝒰_*L,P,K,E*_ and 𝒯_*L,P,K,E*_ is illustrated in Figure 4.

**Fig 4:**
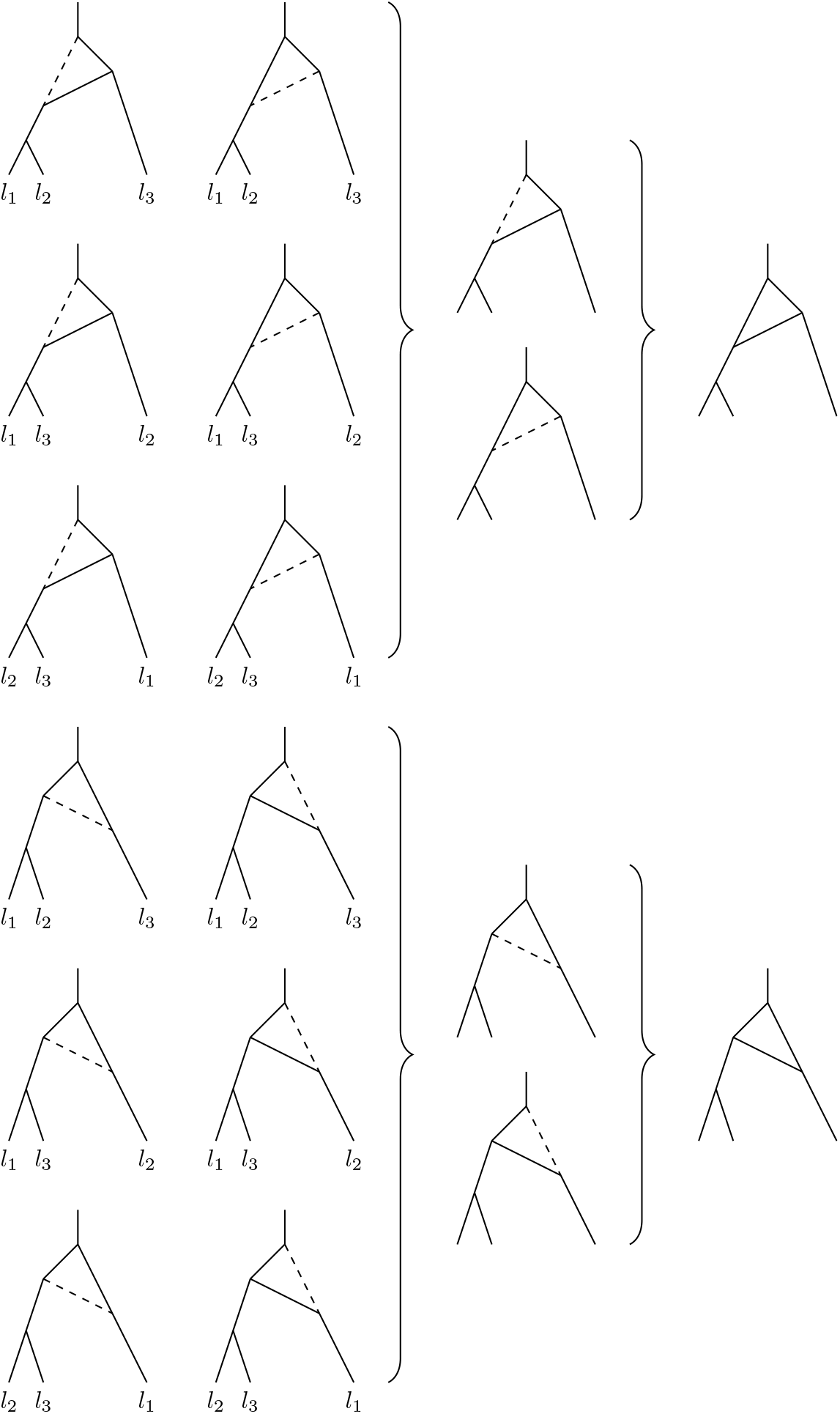
Representatives of the sets 𝒯_3,1,1,0_(left), 𝒰_3,1,1,0_(center) and 𝒮_3,1,1,0_(right). In all of our illustrations on labeled or unlabeled topologies, the admixture edges in *ε*_*A*_ are marked with a dashed line. Here |𝒮_3,1,1,0_| = 2, |𝒰_3,1,1,0_| = 4 and | 𝒯_3,1,1,0_| = *N*(3, 1, 1, 0) = 12.

In Figure 4, both shapes in 𝒮_3,1,1,0_ correspond to two unlabeled topologies in 𝒰_3,1,1,0_, and each of the four unlabeled topologies in 𝒰_3,1,1,0_ correspond to three topologies in 𝒯_3,1,1,0_. However, in general some graphs exhibit more symmetry than others. Let 𝒰_*S*_ be the set of unlabeled topologies corresponding to the shape *S*, and 𝒯_*U*_ the set of topologies corresponding to the unlabeled topology *U*, so that

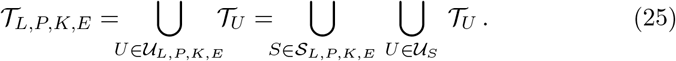

As illustrated in Figure 5, we can have 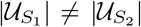 with *S*_1_, *S*_2_ ∈ S_*L,P,K,E*_, and 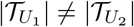 with *U*_1_, *U*_2_ ∈ 𝒰_*S*_, *S* ∈ *S*_*L,P,K,E*_.

**Fig 5:**
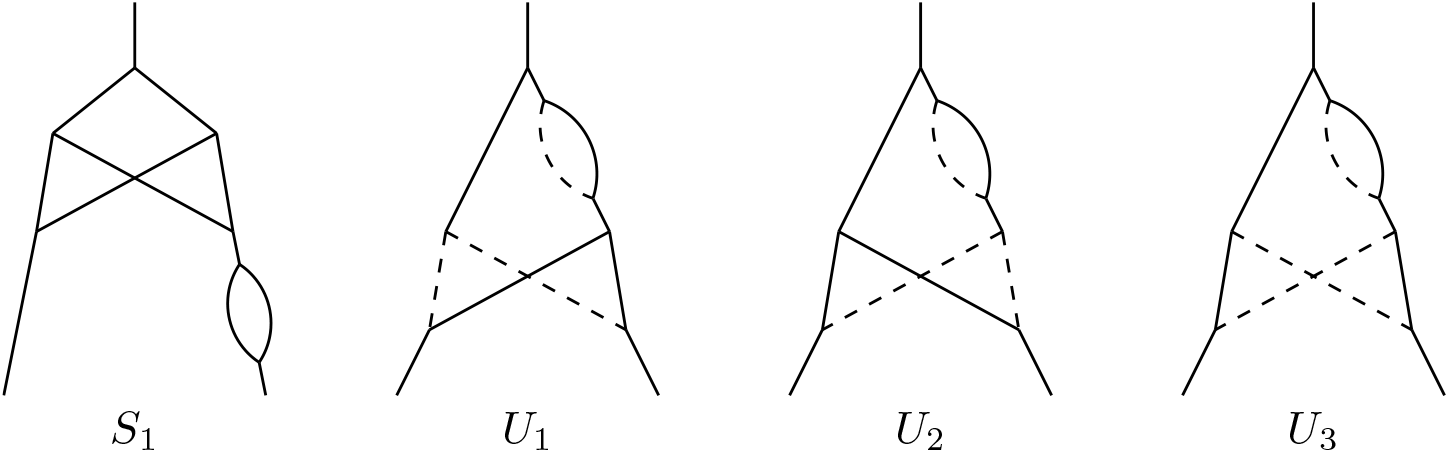
Illustration of a shape, *S*_1_ (left), and the three unlabeled topologies corresponding to a shape *S*_2_ (*S*_2_ not explicitly drawn). We have *S*_1_, *S*_2_ ∈ 𝒮_2,0,3,1_, 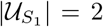 and 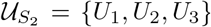. Furthermore, the leaves of *U*_1_ and *U*_2_ are indistinguishable, while the leaves of *U*_3_ can be told apart. To see this, follow the path from the leaves to root; in *U*_1_ and *U*_2_ the path will only depend on whether the parent branch of the first encountered admixture is in *ε*_*M*_ or *ε*_*A*_ and not on the starting leaf. In contrast the starting leaf does matter for *U*_3_ so the leaves are distinguishable. Hence, 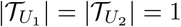 but 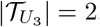.

Given an unlabeled topology *U* ∈ 𝒰_*L,P,K,E*_, choose an arbitrary multigraph topology representative of *U* denoted *G*. Let 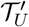 be the set of all multigraph topologies obtained by relabeling the *L* leaves of *G* using the *L*! possible permutations. Clearly each equivalence class in 𝒯_*U*_ is represented by at least one of the elements in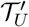, implying 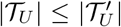. Consider the set of elements of 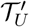 that are isomorphic to *G* under a leaf preserving isomorphism. It can be considered as a set of permutations, *H*_*G*_, where the identity permutation corresponds to *G*. It is straightforward to show that *H*_*G*_ is a subgroup of the permutation group. Because *H*_*G*_ is a subgroup, its cosets are disjoint, contain the same number of elements, and span the whole permutation group (see Figure 6). This characterization gives us a more concrete representation of the elements 𝒯_*L,P,K,E*_, namely as equi-sized sets of permutations of the leaf-labels.

**Fig 6:**
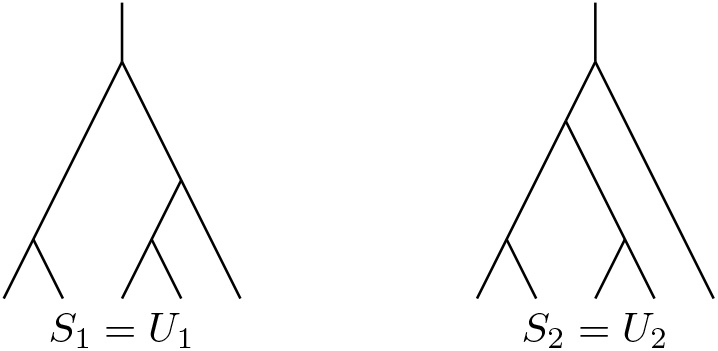
The two shapes of 𝒮(5, 2, 0, 0), denoted *S*_1_ and *S*_2_ are illustrated above. Here, 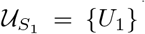 and 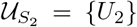, because there are no admixture edges. Interestingly, the shape *S*_2_ exhibits more symmetry than the shape *S*_1_. To see this, let *G*_1_ and *G*_2_ be representatives of *U*_1_ and *U*_2_ with leaves labeled *l*_1_, *l*_2_, *l*_3_, *l*_4_, *l*_5_ from left to right. In both cases 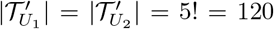. The group 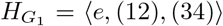 has four elements and so 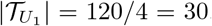. The group 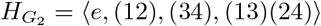 has eight elements and so 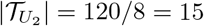. Altogether, *N*(5, 2, 0, 0) = 15 + 30 = 45 by decomposition (25). Notice that the leaves *l*_4_ and *l*_5_ form a pair in one fifth of the elements in both 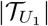 and 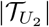 although the two sets are of different size.

There are two basic approaches for counting phylogenetic trees with labeled leaves: recurrence by splitting the tree at the root^33^ or recurrence by removal of one of the leaves^25^. The first approach is difficult to generalize to admixture graphs, but the latter strategy behaves relatively nicely. Our strategy for counting topologies is based on decomposing a topology into a recursive series of predecessors, such that we only need to count the number of possible predecessors in each step. The *predecessor ρ*(*G*) of a labeled topology *G* with *L* leaves is defined as follows. In *ρ*(*G*) the leaf *l*_*L*_ and the terminal edge leading to it are removed and

1) If the terminal edge was from a node with outdegree 2, the edge to it and the remaining edge from it are combined to a single edge.
2) If the terminal edge was from an admixture node, the admixture node is also removed, its parental edge in *ε*_*M*_ is redirected to a new leaf *l_L_* and its parental edge in *ε*_*A*_ is redirected to a new leaf *l*_*L*+1_.

Examples of topologies and their predecessors are given in Figure 7. The topology with only one edge (graph *ρ*(*G*_1.2_) in Figure 7) has no predecessor. By examining the graph elements of the predecessors, we can now derive a recurrence formula for the numbers *N*(*L, P, K, E*):

**Fig 7:**
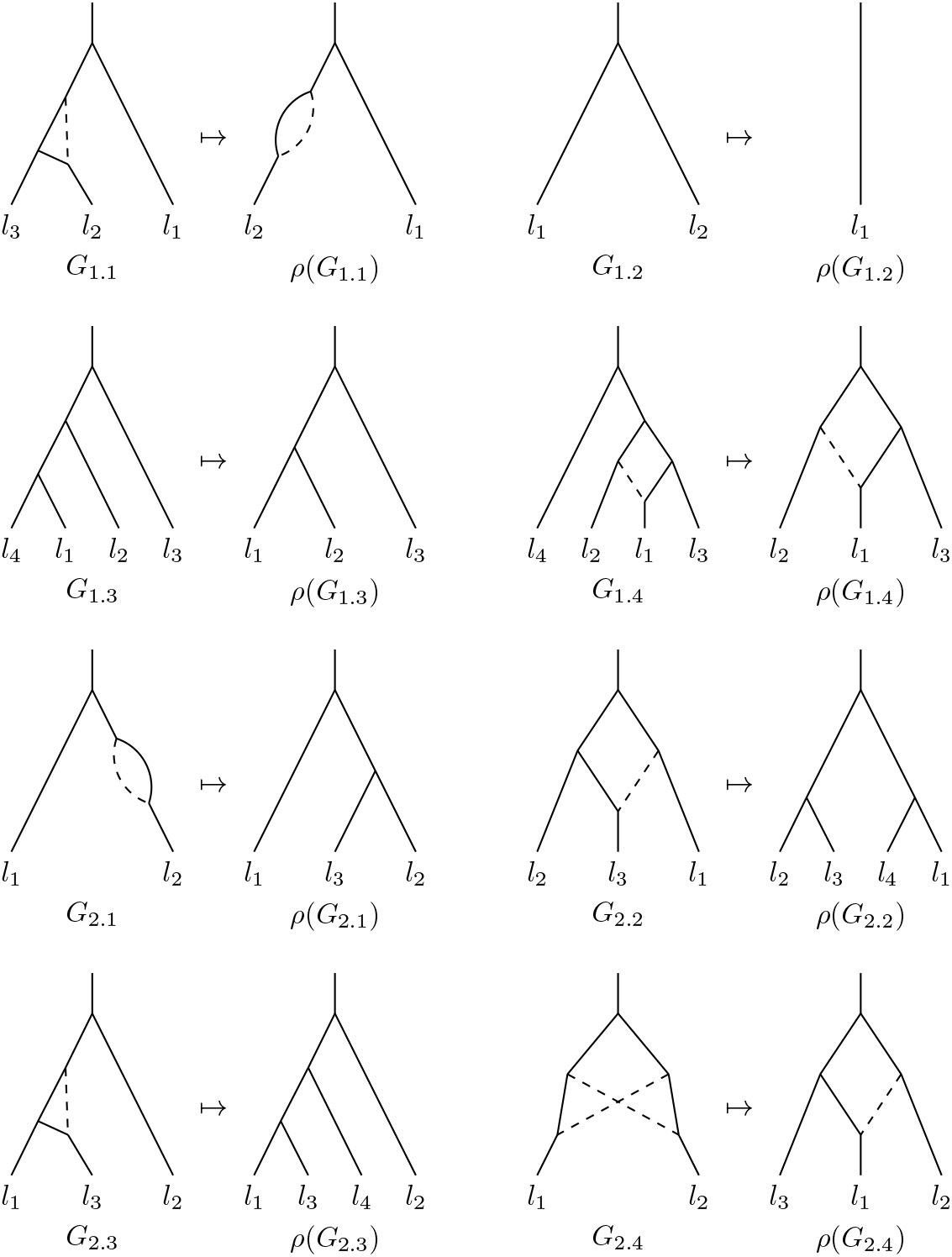
Example graphs and their predecessors from each sub case 1.1) – 2.4). The graph *ρ*(*G*_1.2_) is the only labeled admixture graph that doesn’t have a predecessor, and the ultimate predecessor of every other graph.

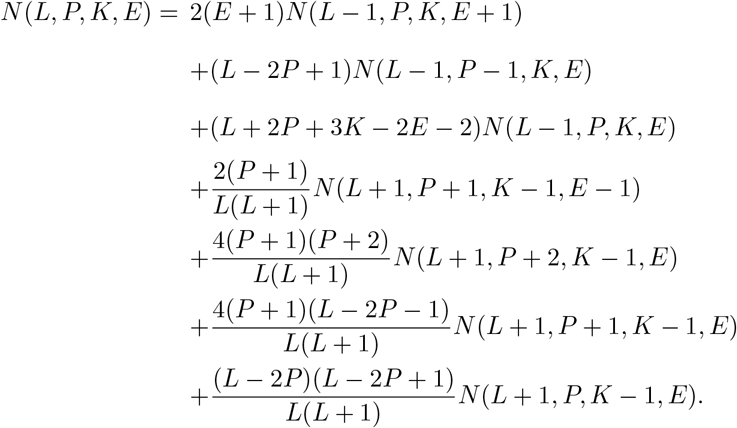

The initial conditions are *N*(1, 0, 0, 0) = 1 and *N*(*L, P, K, E*) = 0 if *L* < 1, *P* > 2*L, K* < *E* or *E* < 0.

The predecessor of any topology in 𝒯_*L,P,K,E*_ is from one of eight possible sources 𝒯_*L′,P′,K′,E′*_. We count *N*(*L, P, K, E*) by looking at these eight sub cases and finding out which graphs in 𝒯_*L′,P′,K′,E′*_ are eligible predecessors and of how many graphs in 𝒯_*L,P,K,E*_. An example of all the sub cases 1.1) – 2.4) is presented in Figure 7.

1.1) The latest leaf *l*_*L*_ stems from an edge forming an eye in *ρ*(*G*). Then *ρ*(*G*) ∈ 𝒯_*L*−1,*P,K,E*+1_, and since every topology in 𝒯_*L*−1,*P,K,E*+1_ has *E* + 1 eyes, and every eye has two edges, the contribution to *N*(*L, P, K, E*) is

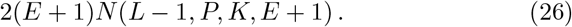
1.2) The latest leaf *l*_*L*_ stems from a terminal edge not belonging to any pairs in *ρ*(*G*). Since *ρ*(*G*) ∈ 𝒯_*L*−1,*P* −1,*K,E*_, and every topology in 𝒯_*L*−1,*P*−1,*K,E*_ has *L* − 1 terminal edges, 2(*P* − 1) of which belong to a pair, the contribution to *N*(*L, P, K, E*) is

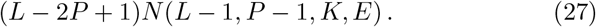
1.3) The latest leaf *l*_*L*_ stems from an edge belonging to a pair in *ρ*(*G*). Since *ρ*(*G*) ∈ 𝒯_*L*−1,*P,K,E*_, and every topology in 𝒯_*L*−1,*P,K,E*_ has 2*P* edges belonging to a pair, the contribution to *N*(*L, P, K, E*) is

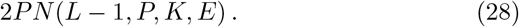
1.4) The latest leaf *l*_*L*_ stems from an edge which is neither terminal nor form an eye in *ρ*(*G*). Since *ρ*(*G*) ∈ 𝒯_*L*−1,*P,K,E*_, and every topology in 𝒯_*L*−1,*P,K,E*_ has 2*L* + 3*K* − 3 edges by induction, *L* − 1 of which are terminal and other 2*E* form eyes, the contribution to *N*(*L, P, K, E*) is

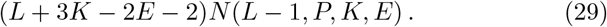
2.1) The latest leaf *l*_*L*_ of *G* stems from an admixture node formed by joining together the edges *l*_*L*_ and *l*_*L*+1_ that form a pair in *ρ*(*G*). We now have *ρ*(*G*) ∈ 𝒯*L*+1,*P* +1,*K*−1,*E*−1, but not every topology in 𝒯*L*+1,*P* +1,*K*−1,*E*−1 have the property 𝔭_1_ that the leaves *l*_*L*_ and *l*_*L*+1_ form a pair. Let *U* ∈ 𝒰_*L*+1,*P*+1,*K*−1,*E*−1_ be any unlabeled topology. By simple combinatorics, the proportion of multigraph topologies with property 𝔭_1_ among the (*L* + 1)! elements in 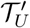 is 2(*P* + 1)*/*(*L*^2^ + *L*). Since the property 𝔭_1_ is invariant under leaf preserving graph isomorphisms, and every equivalence class under the leaf preserving graph isomorphisms in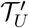 have the same cardinality, the proportion of 𝔭_1_ among the labeled admixture graphs in 𝒯_*U*_ is also 2(*P* + 1)*/*(*L*^2^ + *L*). Finally, because this applies to every *U* ∈ 𝒰_*L*+1,*P* +1,*K*−1,*E*−1_, using (25) we conclude that the proportion of topologies having property 𝔭_1_ among all the topologies in 𝒯_*L*+1,*P*+1,*K*−1,*E*−1_ must be 2(*P* + 1)*/*(*L*^2^ + *L*) too. Therefore, the contribution to *N*(*L, P, K, E*) is

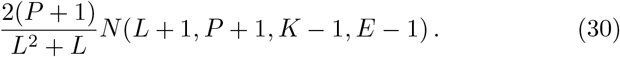
2.2) The latest leaf *l*_*L*_ stems from an admixture node formed by joining together two edges belonging to two distinct pairs in *ρ*(*G*). We now have *ρ*(*G*) ∈ 𝒯_*L*+1,*P* +2,*K*−1,*E*_, but not every topology in 𝒯_*L*+1,*P*+2,*K*−1,*E*_ have the property 𝔭_2_ that the leaves *l*_*L*_ and *l*_*L*+1_ belong to two distinct pairs. Let *U* ∈ 𝒰_*L*+1,*P*+2,*K*−1,*E*_ be any unlabeled topology. By simple combinatorics, the proportion of multigraph topologies having property p_2_ among the (*L* + 1)! elements in 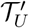 is 4(*P*^2^ + 3*P* + 2)*/*(*L*^2^ + *L*). As before, since the property 𝔭_2_ is invariant under leaf preserving graph isomorphisms, all equivalence classes in 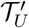 are of equal size and this holds for all unlabeled topologies, the proportion of topologies having property 𝔭_2_ among the elements in 𝒯_*L*+1,*P*+2,*K*−1,*E*_ is the same. Therefore, the contribution to *N*(*L, P, K, E*) is

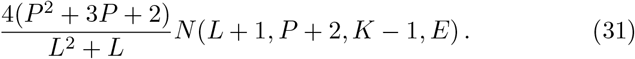
2.3) The latest leaf *l*_*L*_ stems from an admixture node formed by joining together two terminal edges exactly one of which belongs to a pair in *ρ*(*G*). We now have *ρ*(*G*) ∈ 𝒯_*L*+1,*P*+1,*K*−1,*E*_, but not every topology in 𝒯_*L*+1,*P*+1,*K*−1,*E*_ have the property 𝔭_3_ that exactly one of the leaves *l*_*L*_ and *l*_*L*+1_ belong to a pair. Let *U* ∈ 𝒰_*L*+1,*P*+1,*K*−1,*E*_ be any unlabeled topology. By simple combinatorics, the proportion of multigraph topologies having property 𝔭_3_ among the (*L* + 1)! elements in 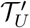 is 4(*PL* + *L* − 2*P*^2^ − 3*P* − 1)*/*(*L*^2^ + *L*). As before, since the property 𝔭_3_ is invariant under leaf preserving graph isomorphisms, all equivalence classes in 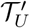 are of equal size and this holds for all unlabeled topologies, the proportion of topologies with property 𝔭_3_ among the topologies in 𝒯_*L*+1,*P*+1,*K*−1,*E*_ is the same. Therefore, the contribution to *N*(*L, P, K, E*) is

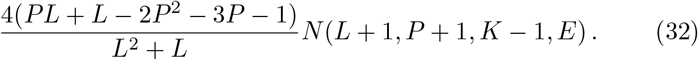
2.4) The latest leaf *l*_*L*_ stems from an admixture node formed by joining together two terminal edges outside the pairs of *ρ*(*G*). We now have *ρ*(*G*) ∈ 𝒯_*L*+1,*P,K*−1,*E*_, but not every topology in 𝒯_*L*+1,*P,K*−1,*E*_ have the property 𝔭_4_ that the leaves *l*_*L*_ and *l*_*L*+1_ do not belong to a pair. Let *U* ∈ 𝒰_*L*+1,*P,K*−1,*E*_ be any unlabeled topology. By simple combinatorics, the proportion of multigraph topologies having property 𝔭_4_ among the (*L* + 1)! elements in 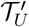 is (*L*^2^ − 4*PL* + *L* + 4*P*^2^ − 2*P*)*/*(*L*^2^ + *L*). As before, since the property 𝔭_4_ is invariant under leaf preserving graph isomorphisms, all equivalence classes in 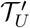 are of equal size and this holds for all unlabeled topologies, the proportion of topologies having property 𝔭_4_ among the elements in 𝒯_*L*+1,*P,K*−1,*E*_ is the same. Therefore, the contribution to *N*(*L, P, K, E*) is

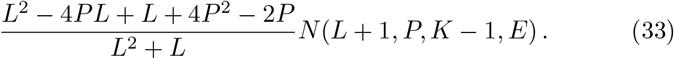

Formula (A.3) follows by summing up all the contributions (26) – (33). The recurrence procedure converges in *L* + 2*K* steps, because either *L* decreases by one, or *K* decreases by one increasing *L* by one.

## B Supplementary Figures

**Fig S1:**
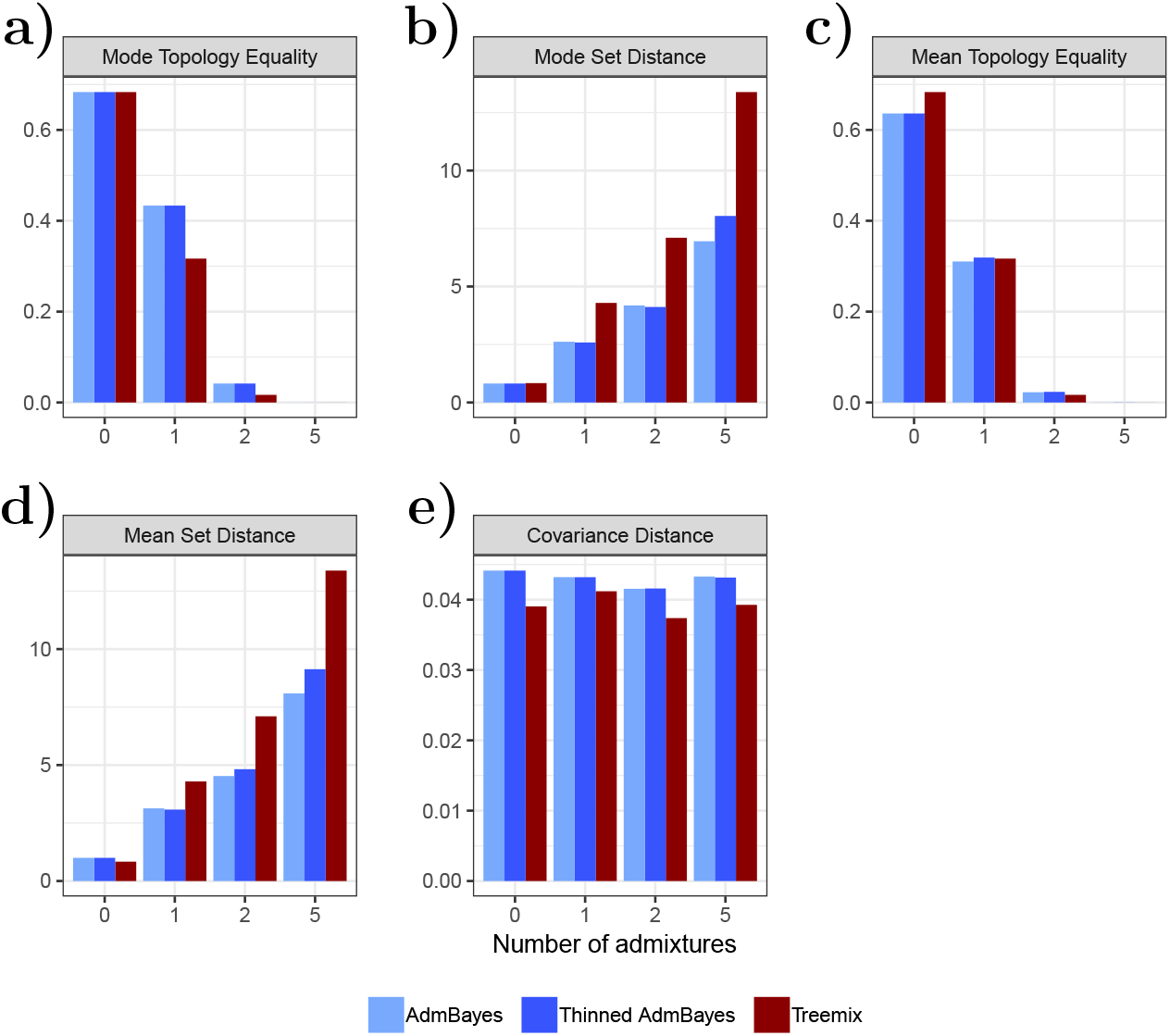
We simulated datasets based on admixture graphs with 10 populations and 0, 1, 2, or 5 admixture events and analyzed them with TreeMix and AdmixtureBayes. For explanation of how we simulated the datasets, see Appendix A.1. The simulations were equally split between datasets with 2, 10, and 50 haplotypes per population and equally split between datasets with 2000 segments and 5000 segments of 500 kb linked sites. Therefore, each column in the plot is a mix of datasets with 2, 10 and 50 haplotypes and 2000 segments and 5000 segments. We compared the results of AdmixtureBayes and TreeMix to the true underlying admixture graph using 5 different measures (see Results). The thinned AdmixtureBayes results are extracted from the unthinned AdmixtureBayes by discarding all graphs that do not contain the true number of admixture events.

**Fig S2:**
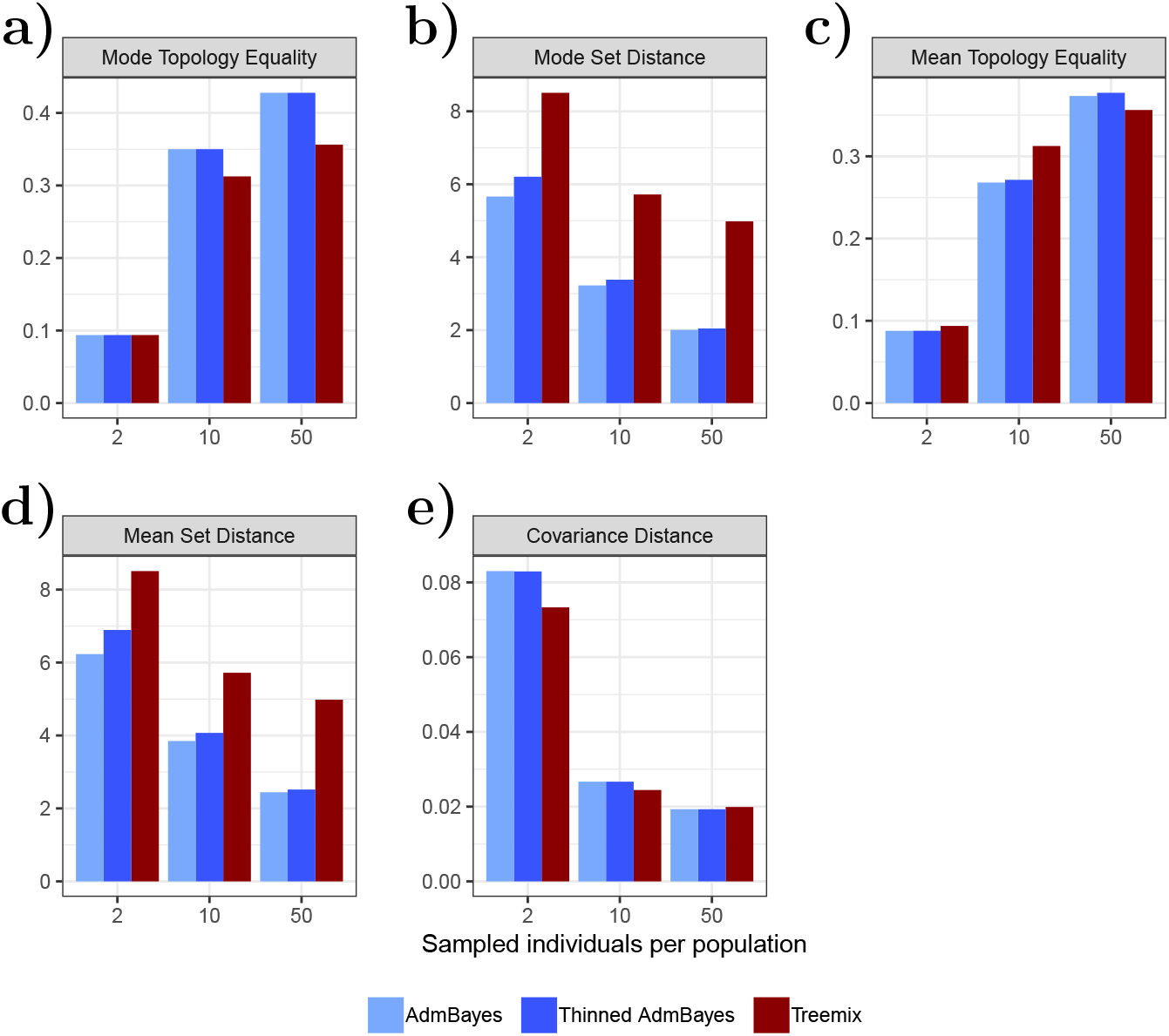
In Figure S1, we separated our simulation study results based on the number of admixture events while averaging over the number of haplotypes per population (2, 10, or 50). These are the same analysis results, instead separated by the number of haplotypes in each population while averaging over numbers of admixture events (0, 1, 2, or 5).

**Fig S3:**
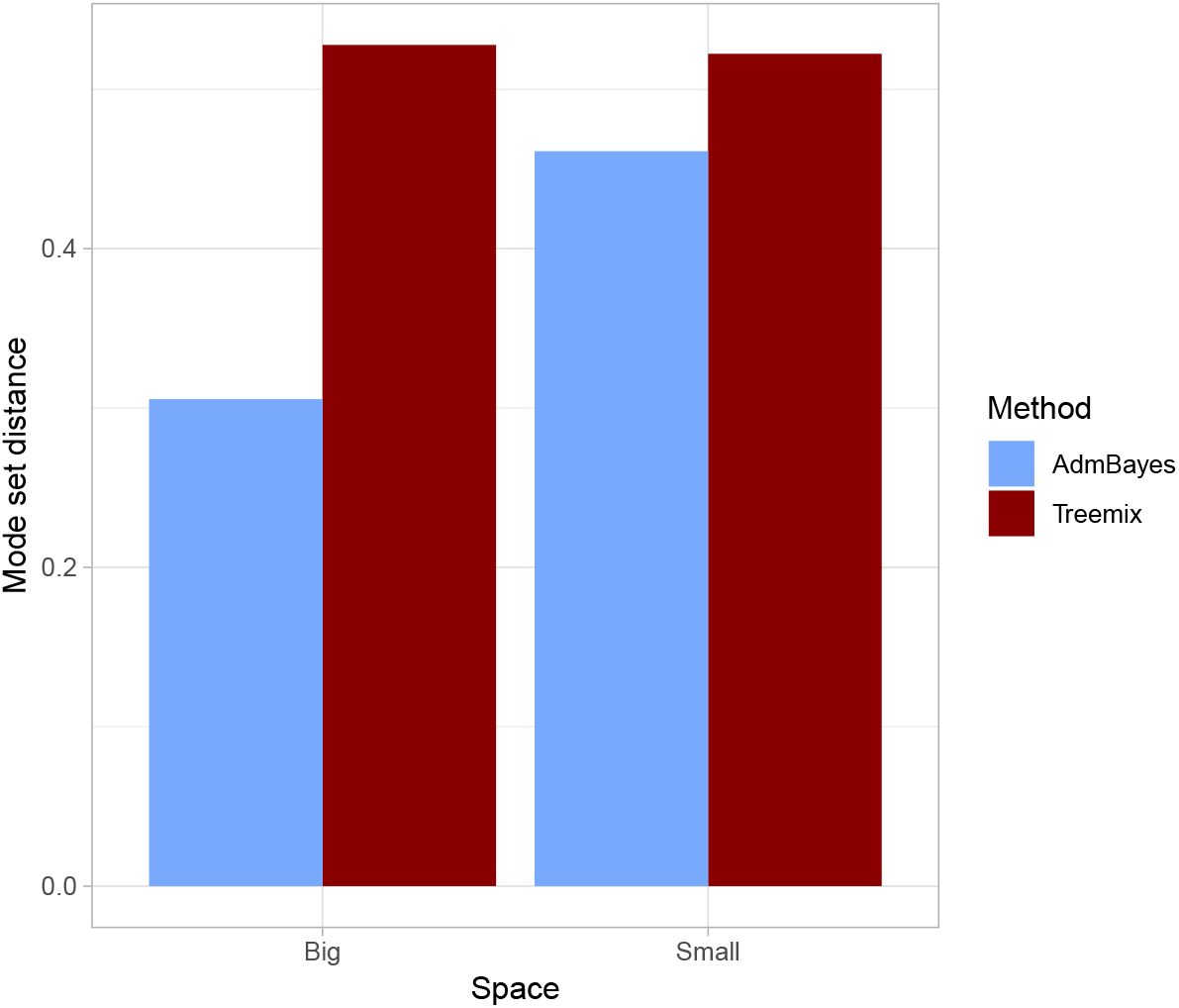
Using admixture graphs with 10 leaves and 0, 1 and 2 admixture events, we simulated 60 different admixture graph datasets with ms (see Appendix A.1). We estimated randomly selected subgraphs of size 3, 4 and 5 from each dataset. The Small column contains graphs built from the marginal dataset and the Big column contains subgraphs of graphs obtained from the full dataset. The Mode Set Distance on the y-axis measures the distance between the true topology of the subgraph and the subgraphs estimated by either AdmixtureBayes or TreeMix.

**Fig S4:**
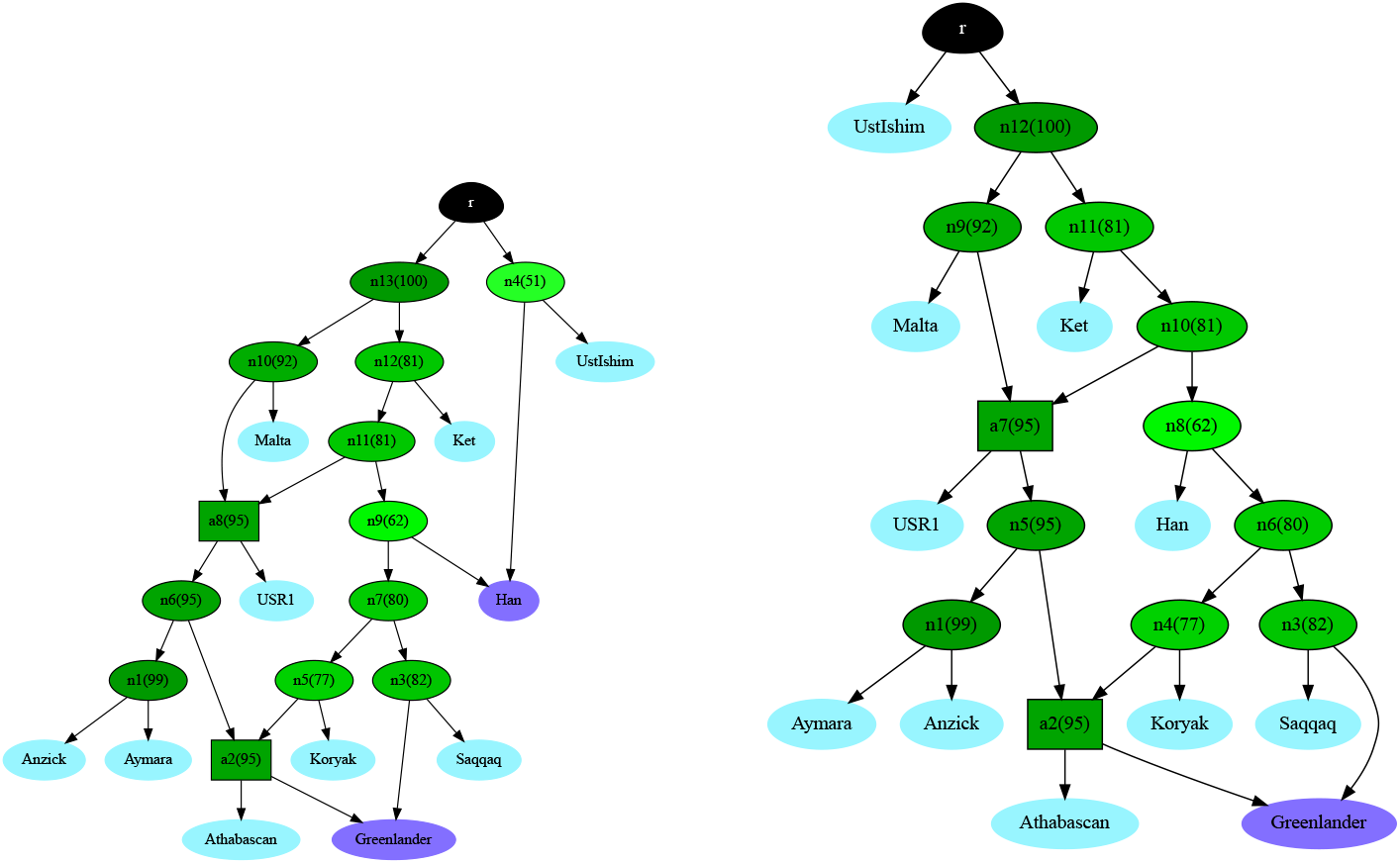
The two minimal topologies with the highest posterior probabilities in our real dataset. Each inner node is colored according to the posterior probability that the true graph has a node with the same descendants. Higher probabilities have a darker shade of green. The posterior probability is written as a percentage in parentheses inside each node, next to the node name, which is arbitrary. The left graph has a posterior of 32%. The right graph has a posterior of 19%.

**Fig S5:**
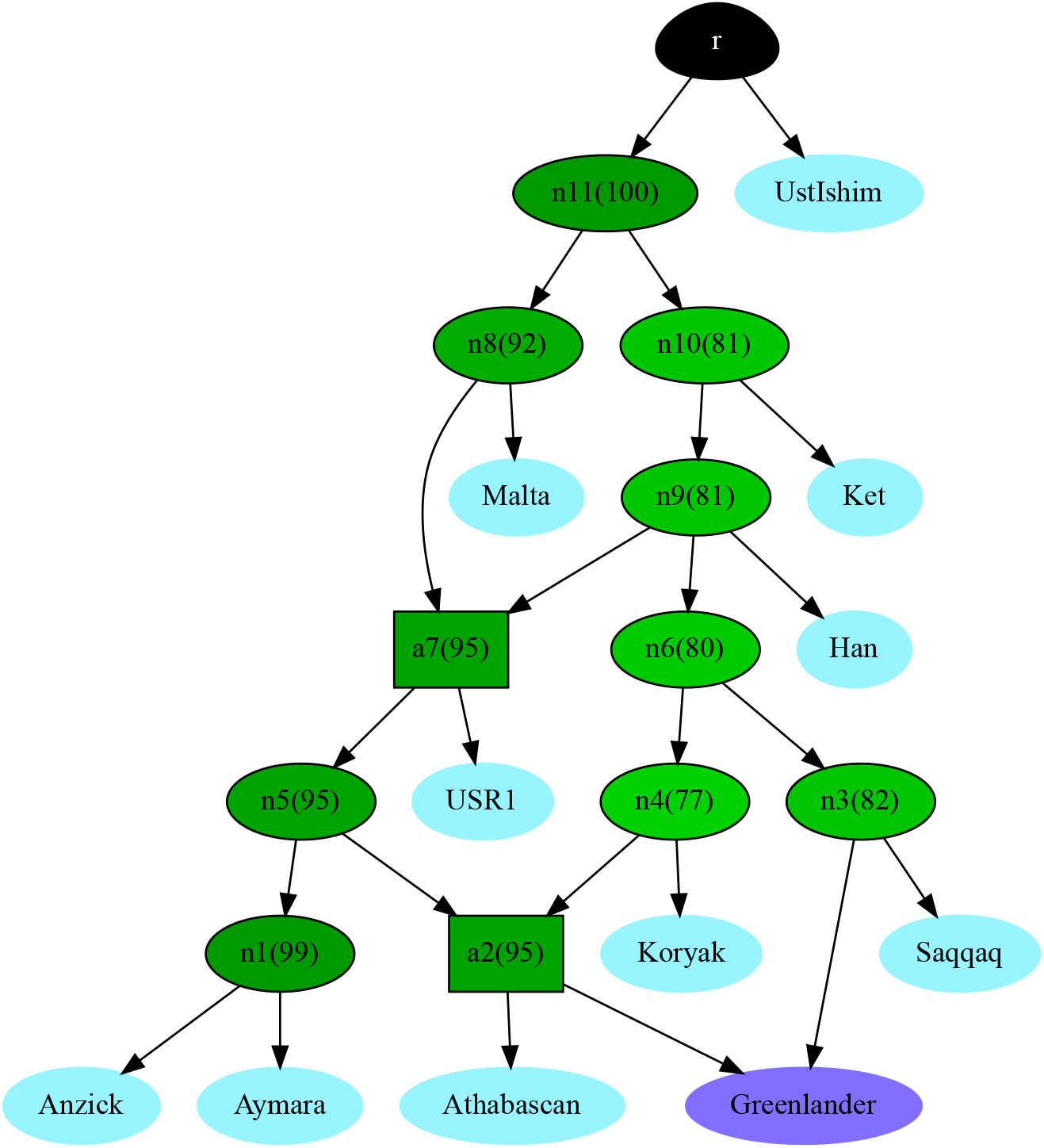
From the posterior AdmixtureBayes samples, we computed the posterior probability of all nodes. The above graph is the smallest directed graph with all the nodes that have a posterior probability higher than 75%. Each internal node is colored according to its posterior probability, as described in Figure S4.

**Fig S6:**
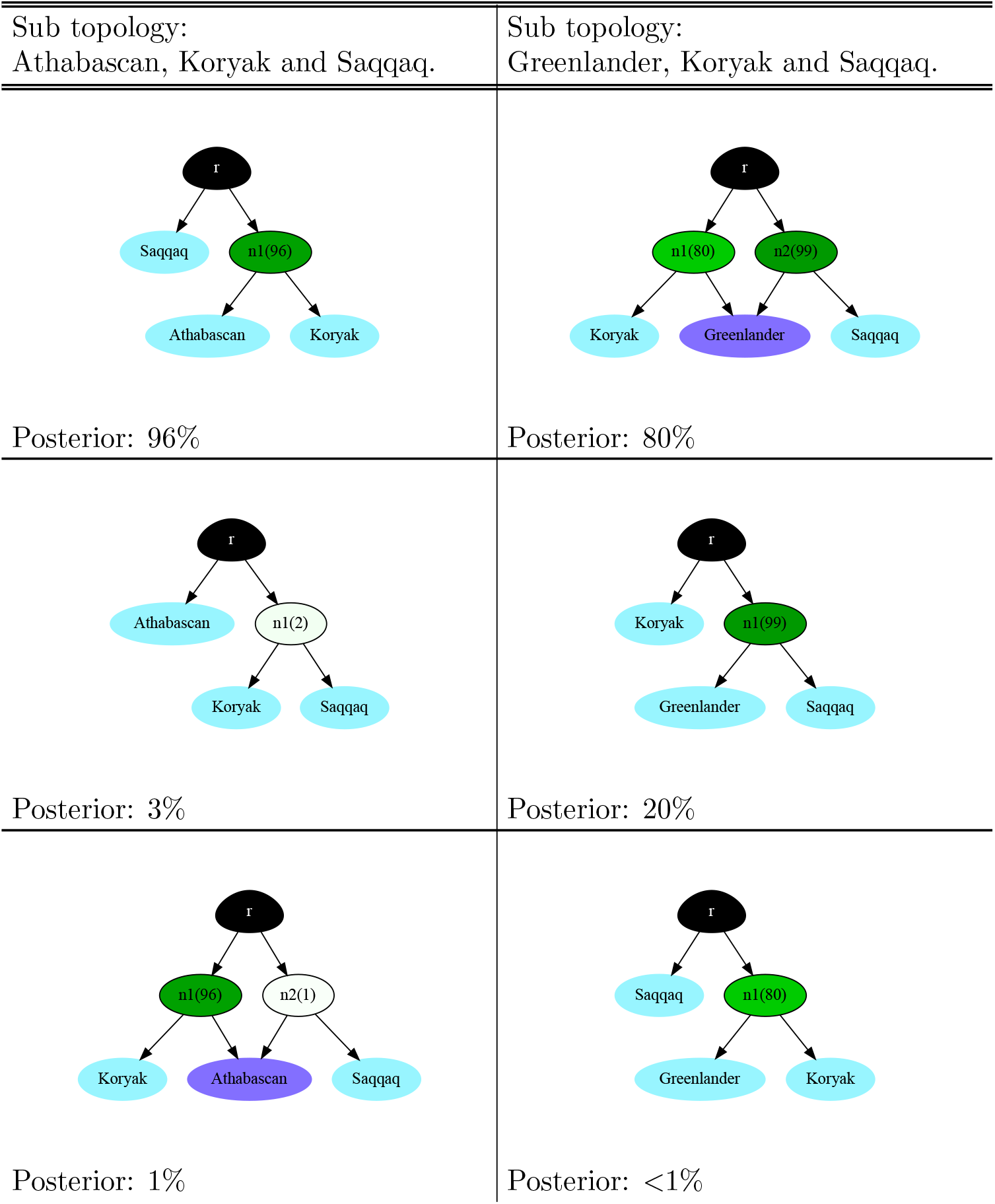
From the posterior AdmixtureBayes sample, we computed the posterior probability of all minimal topologies for several subsets of the populations. Here we show the three topologies with the highest posterior.

**Fig S7:**
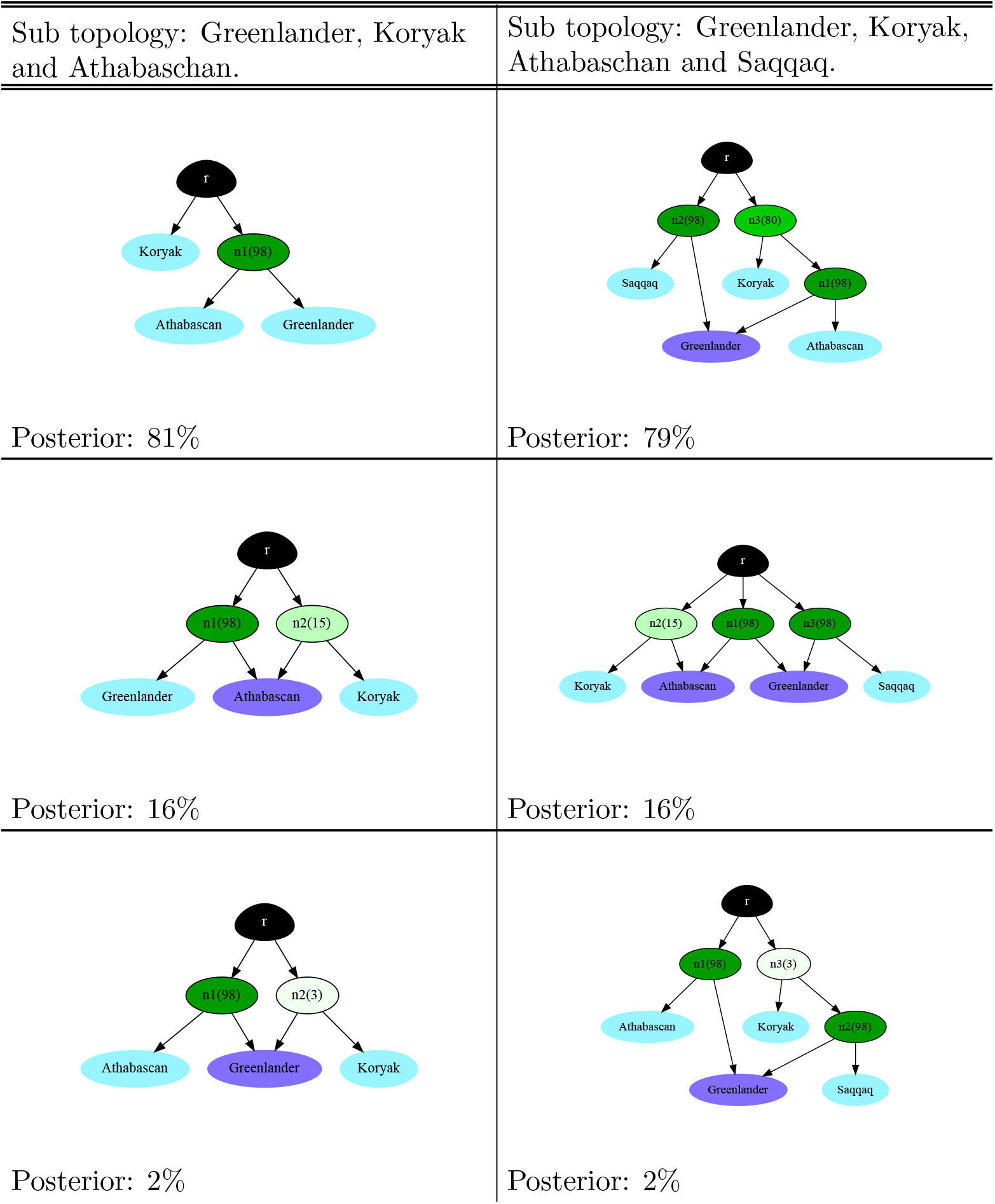
Continuation of Figure S6.

**Fig S8:**
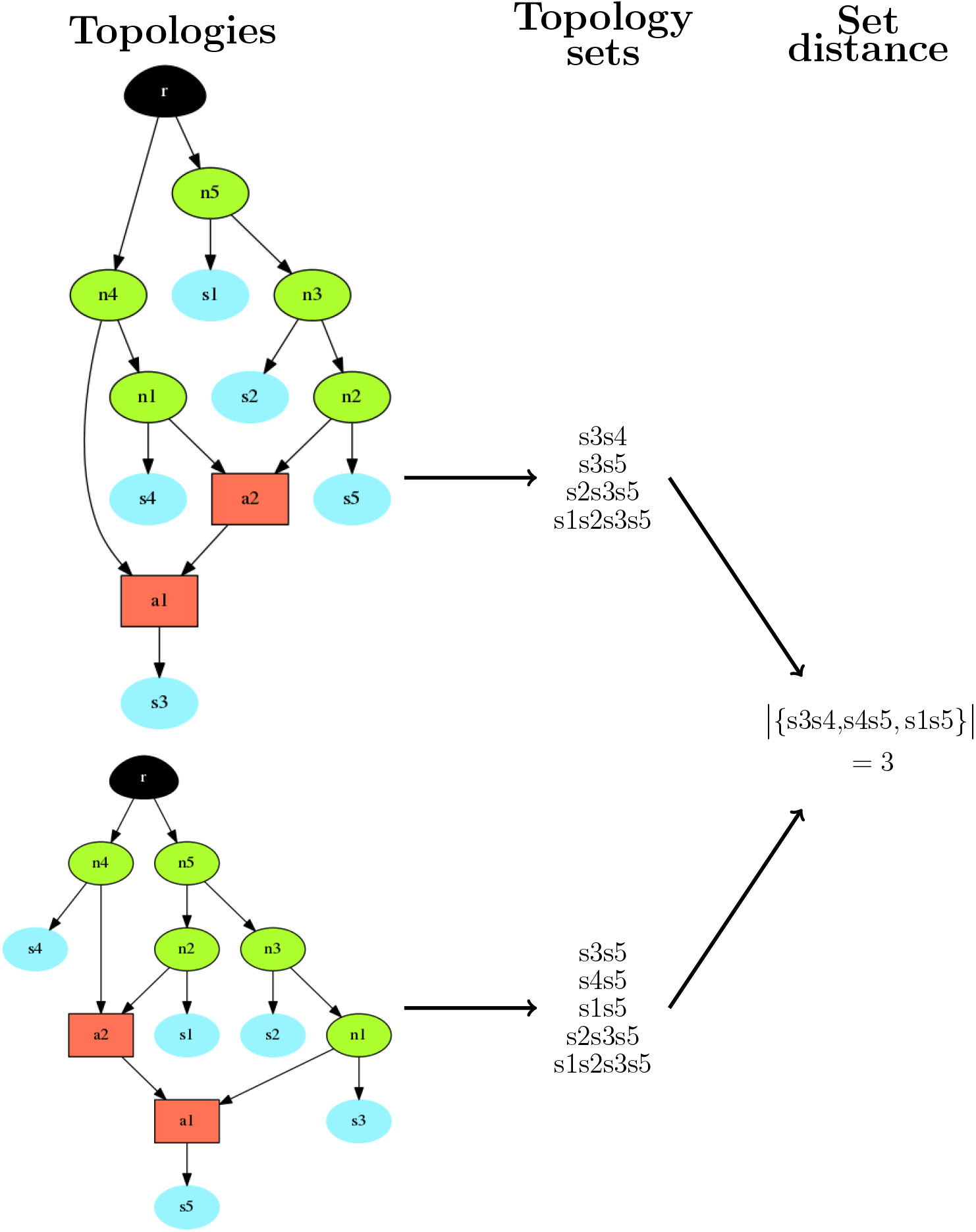
The method used to calculate the Set Distance between two admixture graph topologies (left). First, the topologies are transformed in their descendant sets/topology sets (middle). The distance is then calculated as the symmetric set distance between the two topology sets (right).

**Fig S9:**
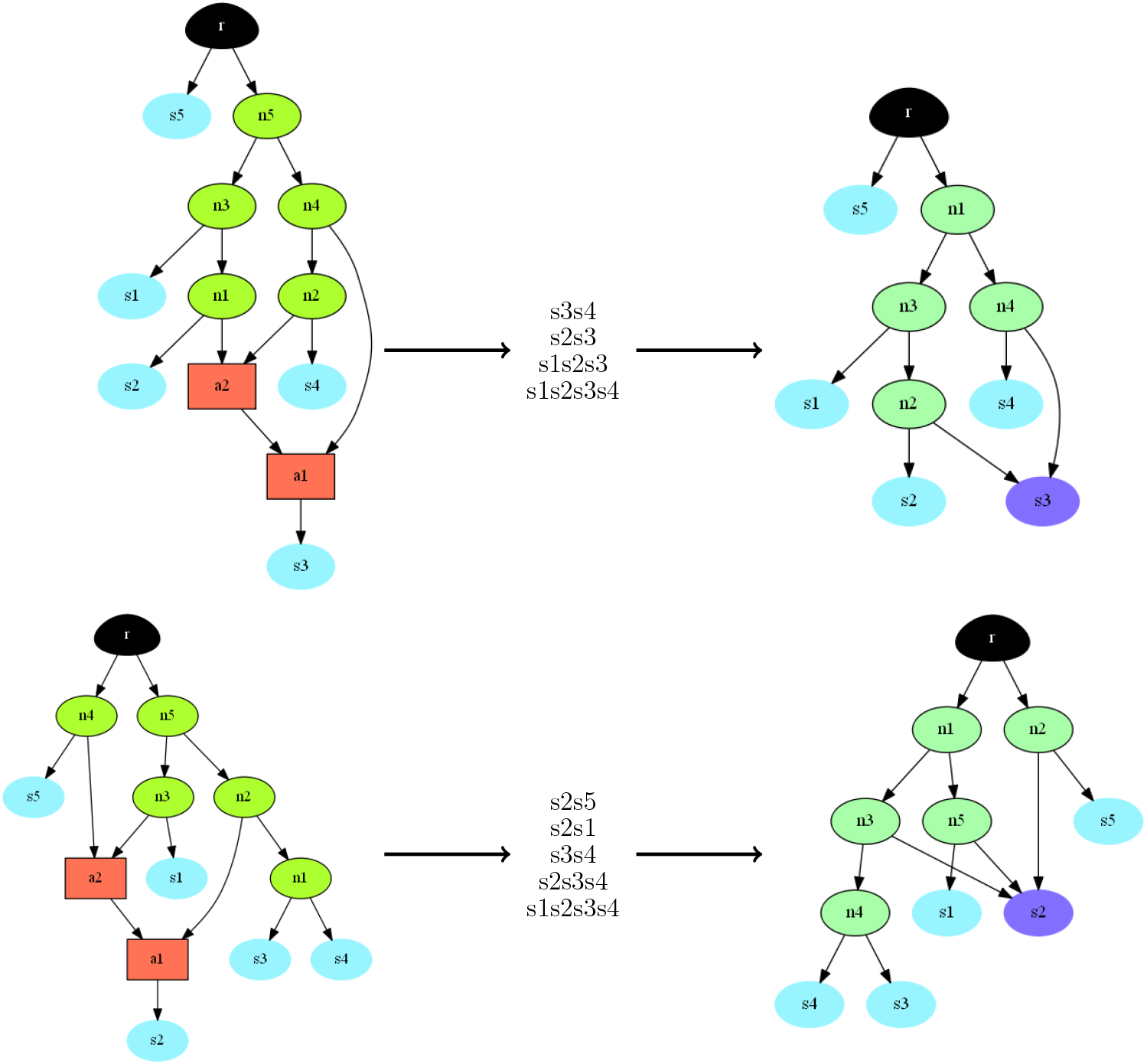
Examples of how the minimal topology is calculated. First, we derive the topology set (middle) from the topology (left). The minimal topology (right) is the smallest possible graph that is consistent with the topology set. Note, node labels assigned to the topology (left) are arbitrary and do not identify corresponding nodes in the minimal topology (right).

**Fig S10:**
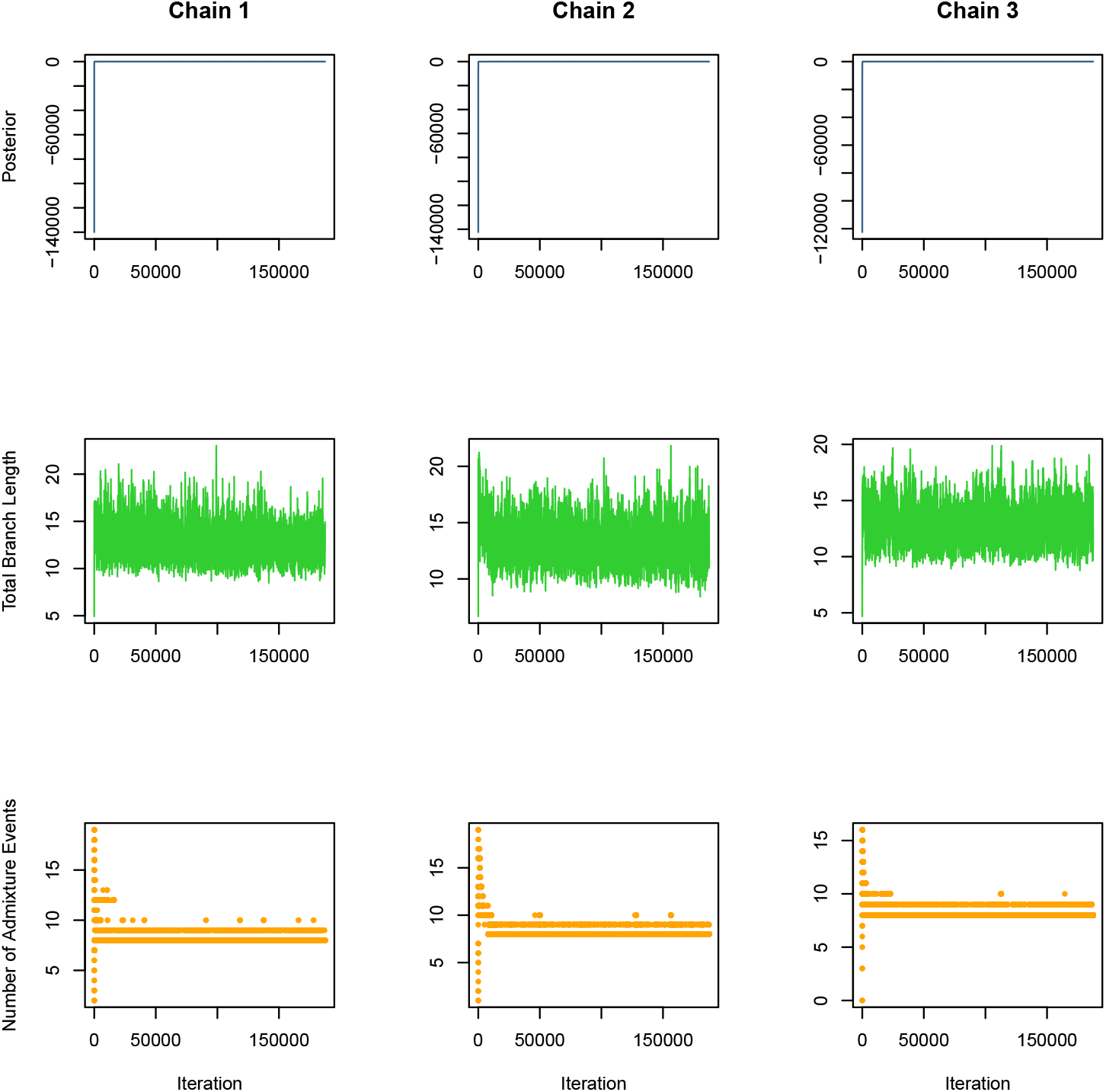
Here, we plot the trace plots for our simulated dataset. Each chain is shown as a separate column. Each summary statistic is shown as a separate row.

**Fig S11:**
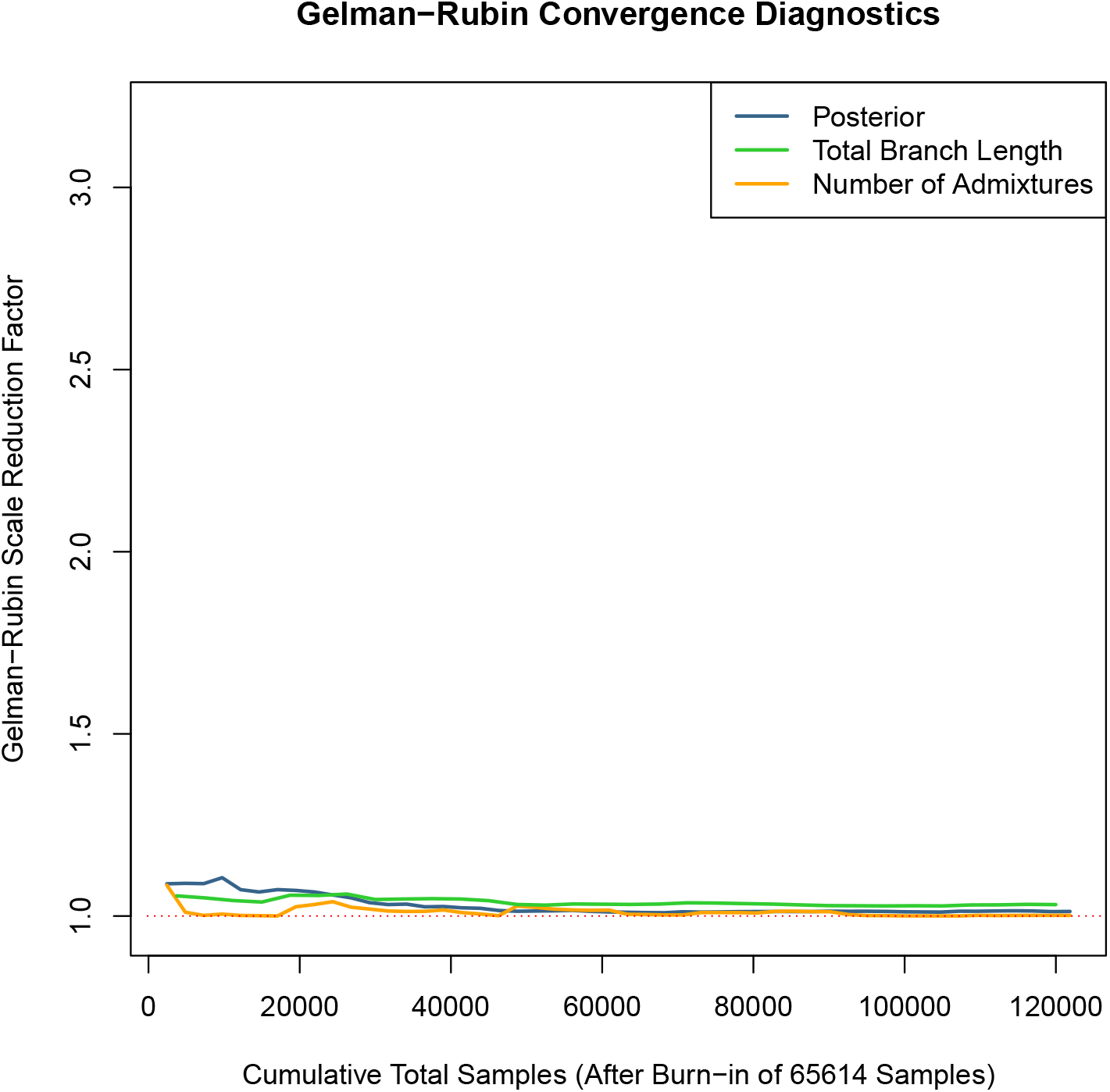
We plot the Gelman-Rubin convergence diagnostics on our simulated dataset for our three summary statistics after a burn-in fraction of 0.35. A rapid convergence to 1 indicates that this is a sufficient burn-in period.

**Fig S12:**
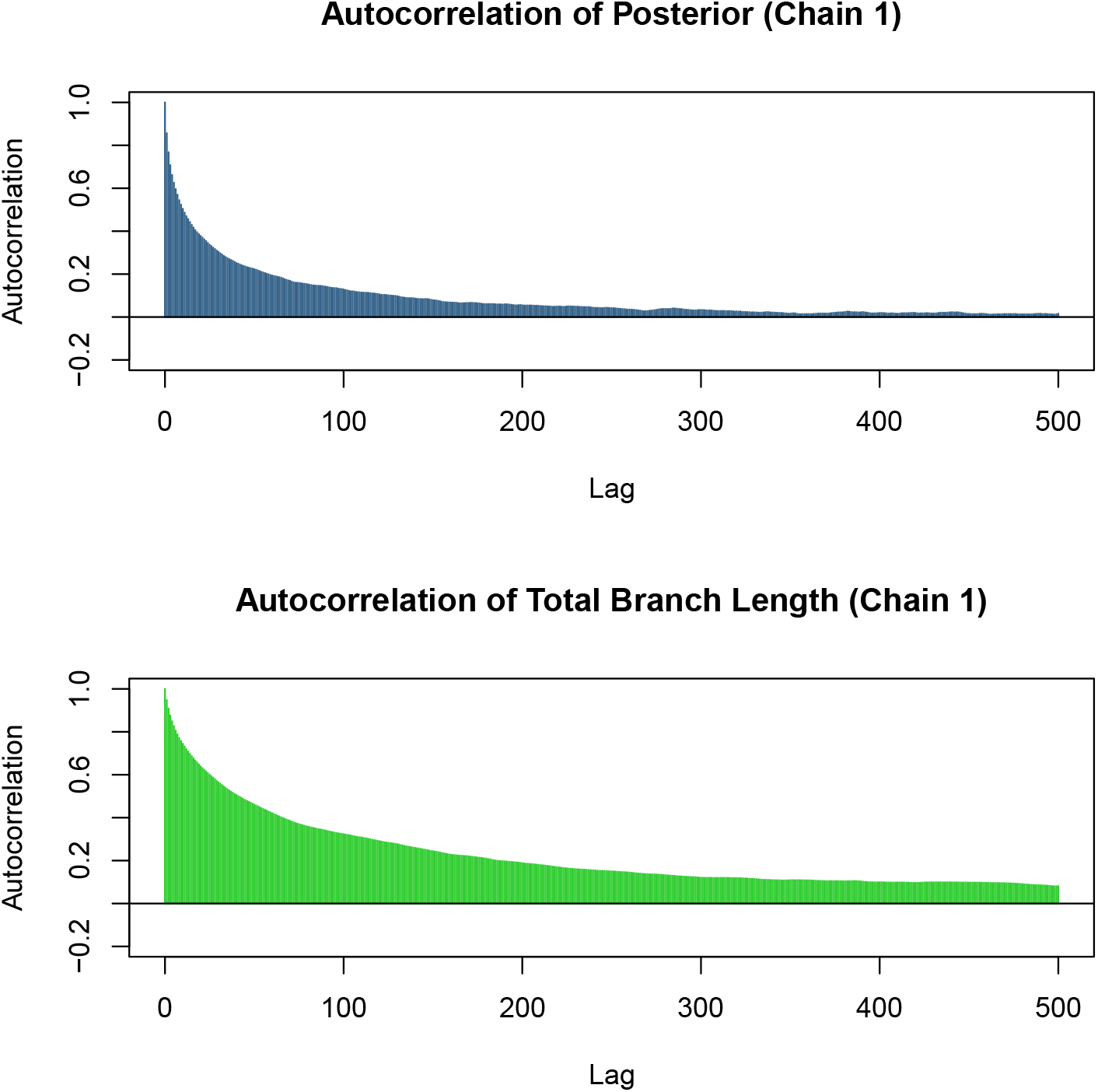
We here show the autocorrelation plots for the summary statistics of our simulated data after a burn-in fraction of 0.35. We only show the results for Chain 1 and do not include the number of admixture events as the autocorrelation shows strange behavior for discrete variables.

**Fig S13:**
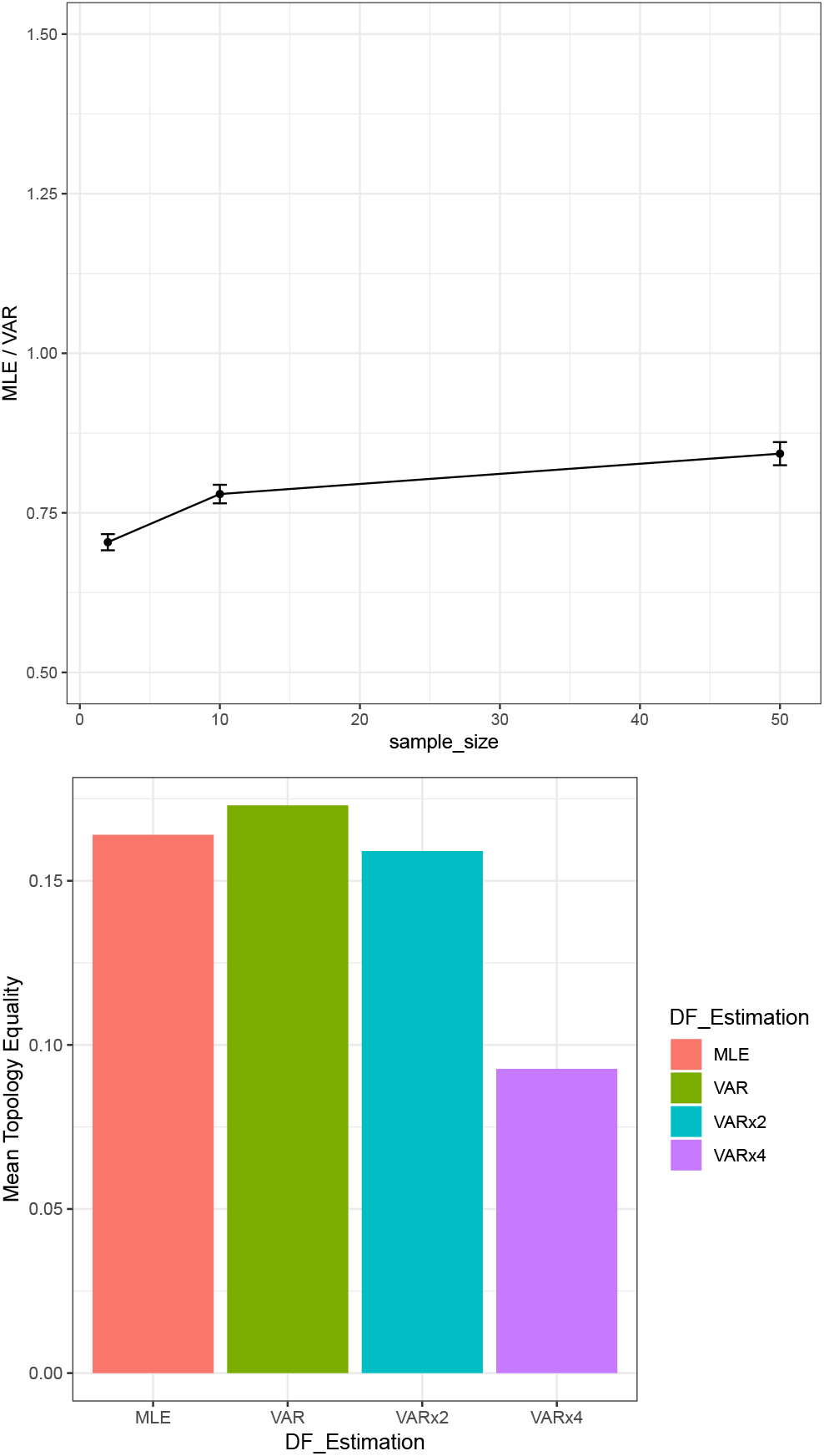
We simulated admixture graphs with 10 leaves and 0, 1 and 2 admixture events. Using these graphs, we simulated datasets using ms with different sample sizes. The top plot illustrates the ratio between the maximum likelihood degrees of freedom estimate from (6) and the variance estimator in (7). We ran AdmixtureBayes with the maximum likelihood estimate (MLE), the variance estimate (VAR), and 2 and 4 times the variance estimate (VARx2 and VARx4 respectively). We calculated the Mean Topology Equality, which was maximized when using the VAR estimates.

**Fig S14:**
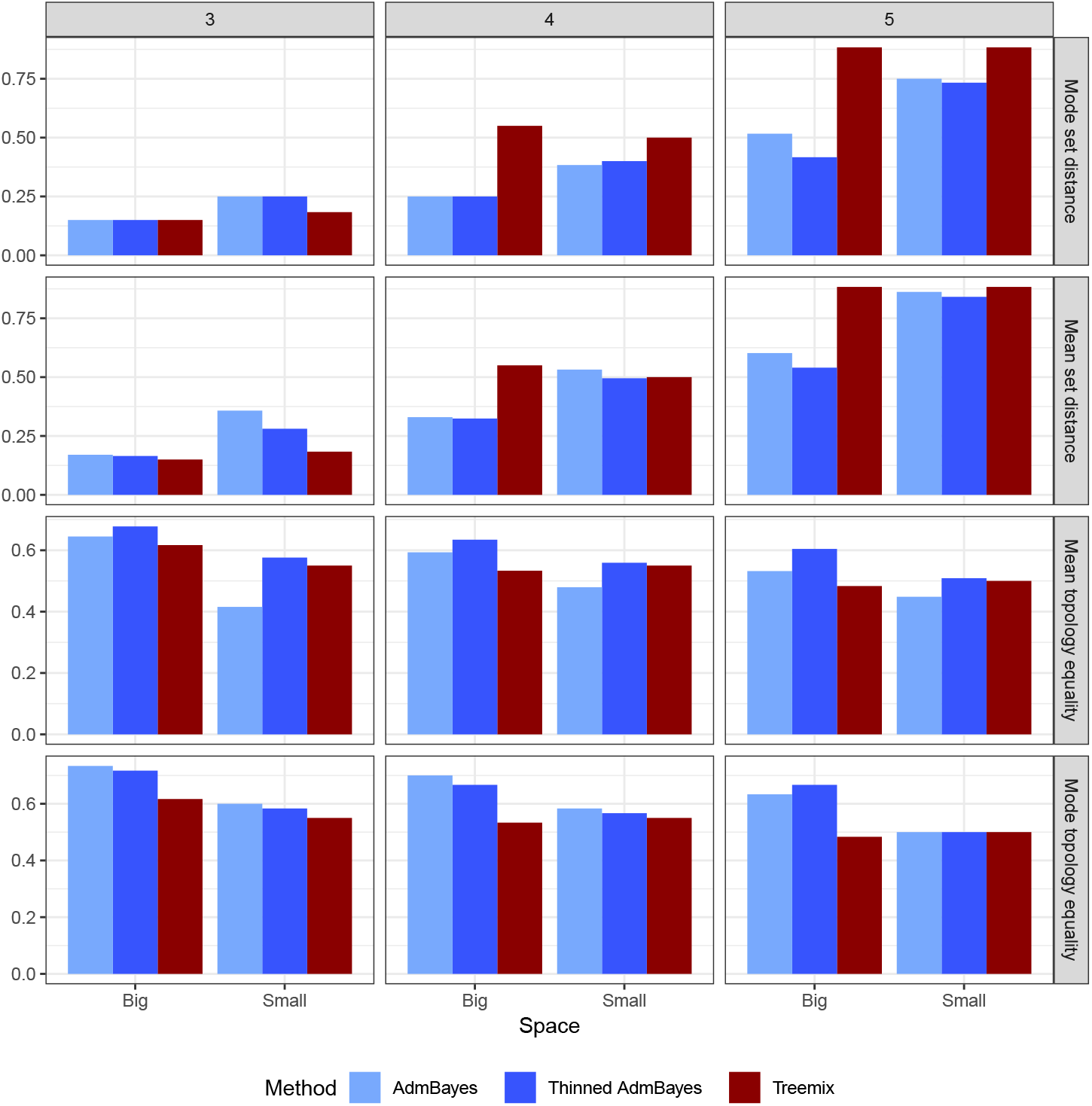
In Figure S3, we calculated how TreeMix and AdmixtureBayes performed when estimating subgraphs. Here we have stratified the same analysis according to subgraph size (in the columns) and measure of accuracy (in the rows)

**Fig S15:**
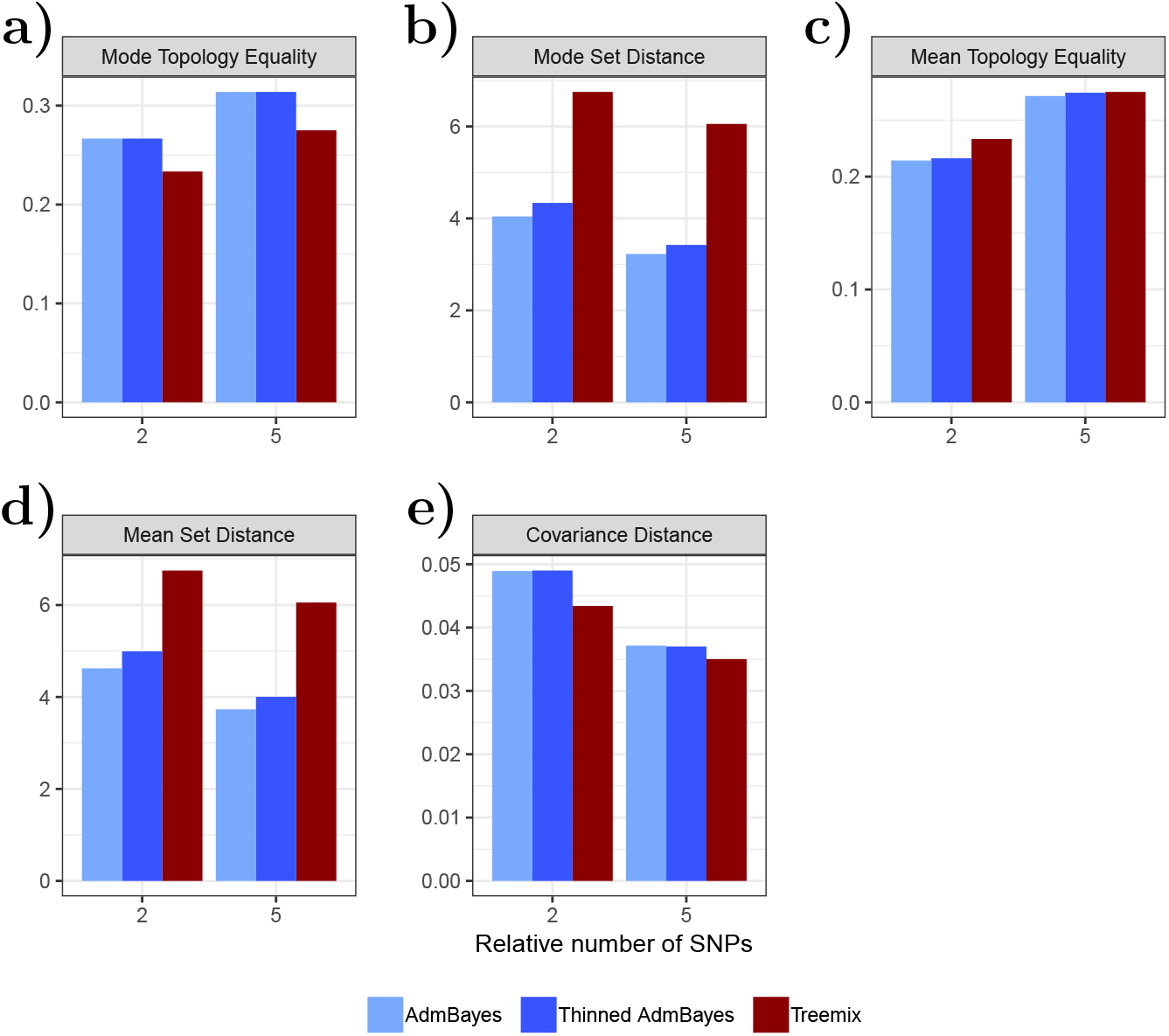
In Figure S1, we separated our simulation study on the number of admixture events. Here, it is separated on the length of the simulated genomes.

## Notes

### Competing Interest Statement

The authors have declared no competing interest.

